# Caspase-14 recognizes and processes IL-1β in epithelial cells to drive anti-bacterial IgG production

**DOI:** 10.64898/2026.04.14.717951

**Authors:** Mingtong Ma, Lin Wang, Baoxue Ge

## Abstract

Caspases-mediated processing of cytokines coordinates cell-autonomous defenses and induction of systemic inflammation ^1^. While caspase-1 processes IL-1β and IL-18 ^2–5^, human caspase-4 processes IL-18 mainly in monocytes ^6^. Caspase-14 is an exception, specializing in epidermal differentiation^7,8^, yet no cytokine target has been firmly established for caspase-14. Here, we report that recognition and IL-1β maturation of IL-1β by caspase-14 in epithelial cells determined anti-bacterial humoral immunity against *Yersina pseudotuberculosis (Y. pseudotuberculosis)* infection. Upon TAK1 inhibition by YopJ, activated caspase-8 cleaved caspase-14 at Asp 146, generating an active 16-kDa fragment, whose exposed pocket directly interacted with and cleaves pro-IL-1β at Cys132. Moreover, conditional knock-out of caspase-14 in epithelial cells or knock-in of a caspase-inactive caspase-14^C136A^ mutant impaired *Y. pseudotuberculosis* induced IL-1β production and eliminated the total anti-*Y. pseudotuberculosis* IgG production, leading to uncontrolled *Y. pseudotuberculosis* infection. Thus, our findings establish caspase-14 as a processor of IL-1β in epithelial cells to propel anti-bacterial humoral immunity, providing insights into the inflammation and vaccine development.

## Main

Inflammasomes coordinate induction of systemic inflammation and host immune defenses ^1,9,10^. In the canonical inflammasome, a nucleotide-binding-oligomerization-domain-like receptor (NLR) senses pathogen products or endogenous dangers to activate caspase-1, which cleaves pro-IL-1β into active forms ^11^. The noncanonical inflammasome involves activation of mouse caspase-11 or its human orthologues caspases-4/5 ^12,13^. Activated human caspase-4, but not mouse caspase-11, directly and efficiently processes IL-18 *in vitro* and during bacterial infections^6^. Activated caspase-1 or caspase-4/5/11 cleaves GSDMD to liberate the pore-forming GSDMD-N domain from the inhibitory GSDMD-C domain ^12,13^. The GSDMD-N domain forms large pores (20–25 nm) on the plasma membrane to trigger pyroptosis, enabling the release of mature IL-1β or IL-18 ^14–18^. Pyroptosis can clear intracellular pathogens ^19,20^ and knock-out of GSDMD impairs humoral immunity in bacteria-infected mice^6^. However, the mechanisms underlying the relation of IL-1β maturation to humoral immunity remain unknown.

Alternatively, macrophage infection by the pathogenic bacteria *Yersinia* or mimic stimulation of lipopolysaccharide (LPS) and transforming growth factor-β-activated kinase 1 (TAK1) specific inhibitor 5z-7-oxozeaenol (5z7) or tumor necrosis factor (TNF) and TAK1 inhibitor induces caspase-8-mediated gasdermin D (GSDMD) cleavage and pyroptosis. During pathogenic *Y. pseudotuberculosis* infection, the *Y. pseudotuberculosis* effector protein YopJ inhibits the activation of TAK1, which is critical for host inflammatory and pro-survival signaling ^21–23^. Inhibition of TAK1 by YopJ leads to an alternate pyroptotic pathway mediated by Toll-like receptors (TLRs) or death receptors that form a complex with the adaptor Fas-associated death domain (FADD) and receptor-interacting serine-threonine protein kinase 1 (RIPK1) and caspase-8. This results in the phosphorylation of RIPK1 to drive the activation of caspase-8, and the cleavage of downstream caspases-1, −3, −7, −9, and −11 ^24,25^. The activated caspase-8 cleaves GSDMD at Asp276 to trigger pyroptosis ^14–17^.While active caspase-8 can cleave pro-IL-1β *in vitro*, studies using *Ripk3 Casp8* macrophages ^26–28^ or no-cleavable *Casp8^D387A/D387A^* mice ^29^ suggest caspase-8 may not directly process IL-1β *in vivo*, but might act through downstream effectors^30^. Very recently, we reported that serine/threonine-protein kinase RIO2 (RIOK2) interacts with the FADD, and RIOK2’s kinase activity drives the transport of lysosome to ER through activating myosin II and thereby translocating FADD-RIPK1-caspase-8 complex from lysosome to ER ^18^. Importantly, RIOK2’s ATPase activity enhances its binding to this complex and directly triggers caspase-8 and gasdermin D cleavage both at ER and in vitro. Furthermore, RIOK2-mediated pyroptosis enhances host defense against *Y. pseudotuberculosis* infection^18^.

*Yersinia pestis*, a Gram-negative coccobacillus, is the pathogen responsible for pestis ^31–33^. Pestis is mainly transmitted by the bite of rat fleas, but also by the respiratory tract, digestive tract, damaged skin and mucous membrane ^31,32,34^. The primary pneumonic plague represents the most severe and rapidly progressing manifestation of the disease, which arises from the inhalation of infectious respiratory droplets containing the bacterium ^34–36^. Previous studies have mainly focused on the maturation and function of IL-1β in macrophages infected with *Y. pseudotuberculosis*, the causative agent of the pestis ^16,35^. In addition to macrophages, epithelial cells, which serve as the first line of host defense, can also be directly infected by *Y. pseudotuberculosis* ^11,37–40^. However, the maturation and function of IL-1β in epithelial cells infected by *Y. pseudotuberculosis* remain largely neglected.

Though caspases are classified as inflammatory (caspase-1, -4, -5, -11) ^8,41,42^ or apoptotic ^8,41,42^, caspase-14 is an exception, specializing in epidermal differentiation. *Caspase-14* is highly expressed in keratinocytes or epithelial cells rather than macrophages ^8,43–45^. It has been shown that caspase-14 is processed by chymotrypsin-like serine protease kallikrein-related peptidase-7 (KLK7) to yield the p17/p10 mature form during keratinocyte terminal differentiation ^46^. After being processed during keratinocyte differentiation ^7,47^, caspase-14 cleaves pro-filaggrin into filaggrin, which is essential for the skin barrier, highlighting functional diversity within the protease family. However, whether caspase-14 is involved in the inflammation of epithelial cells remains unknown.

In this study, we found that caspase-14 recognizes and cleaves IL-1β in epithelial cells. In response to *Y. pseudotuberculosis* infection or LPS plus 5z7 stimulation, activated caspase-8 cleaves caspase-14 at the Asp 146 site, resulting in a 16-kDa active fragment. The activated fragment of caspase-14 unlocks the catalytic site and cleaves pro–IL-1β. Conditional deletion of caspase-14 or knock-in of a caspase-inactive caspase-14^C136A^ mutant impaired *Y. pseudotuberculosis* -induced IL-1β production and eliminated the total anti-*Y. pseudotuberculosis* IgG production, leading to uncontrolled *in vivo* growth of the *Y. pseudotuberculosis*, *Pseudomonas aeruginosa* or *Candida albicans* infection.

## Results

### IL-1β from epithelial cells determines early anti-bacterial immunity

Pestis is mainly transmitted by the bite of rat fleas, but also by the respiratory tract, digestive tract, damaged skin and mucous membrane ^31,32,34^. Previous studies have mainly focused on the maturation and function of IL-1β in macrophages infected with *Y. pseudotuberculosis*, the causative agent of the pestis ^16,35^. In addition to macrophages, epithelial cells can also be directly infected by *Yersinia* ^11,37,38^. To compare the production level and function of IL-1β from macrophages versus epithelial cells, we crossed *Il1b^flox/flox^* (*Il1b^f/f^*) mice with Ccsp-Cre or Lyz2*-Cre* transgenic mice. This generated two conditional knockout strains: *Il1b^flox/flox^*; Ccsp-Cre (*Il1b^f/f^ Ccsp^Cre^*) mice that carried a specific deletion of *Il1b* gene in lung epithelial cells ^48,49^ and *Il1b^flox/flox^* ; Lyz2*-Cre* (*Il1b^f/f^ Lyz2^Cre^*) mice, that has conditional knocked out of *Il1b* from macrophages. Subsequently, C57BL/6 *Il1b^f/f^*, C57BL/6 *Il1b^f/f^ Ccsp^Cre^*, C57BL/6 *Il1b^f/f^ Lyz2^Cre^*, wild-type C57BL/6, and C57BL/6 *Il1b^-/-^* mice were aerosol infected with *Y. pseudotuberculosis* at approximately 200 CFU per mouse. Conditional deletion of *Il1b* from epithelial cells, but not from macrophages almost eliminated the production level of IL-1β from *Y. pseudotuberculosis*-infected mice lung tissues as the whole knock-out of IL-1β from the mice did during the first four days post-infection (**Extended Data Fig.1a)**, suggesting epithelial cells as the main producer of IL-1β at the early stage of infection.

Even after 6 days infection, knock-out of the IL-1β from epithelial cells impaired the lung tissues IL-1β production as those IL-1β knock-out macrophages did (**Extended Data Fig.1a)**. Consistently, the lung tissues of C57BL/6 *Il1b^f/f^ Ccsp^Cre^*mice had the increased bacterial burden comparable as those of *Il1b^-/-^* mice after 3 days infection (**Fig. 1a**), suggesting a decisive role of IL-1β from epithelial cells in the control of bacteria clearance *in vivo*. Even after 6 days of infection, the increased bacterial burden in the lung tissues of C57BL/6 *Il1b^f/f^ Ccsp^Cre^* mice was similar to those of C57BL/6 *Il1b^f/f^ Lyz2^Cre^* mice (**Fig. 1a**). The results suggest that IL-1β from epithelial cells may determine early inflammation and host defense against *Y. pseudotuberculosis* infection.

**Fig. 1.**
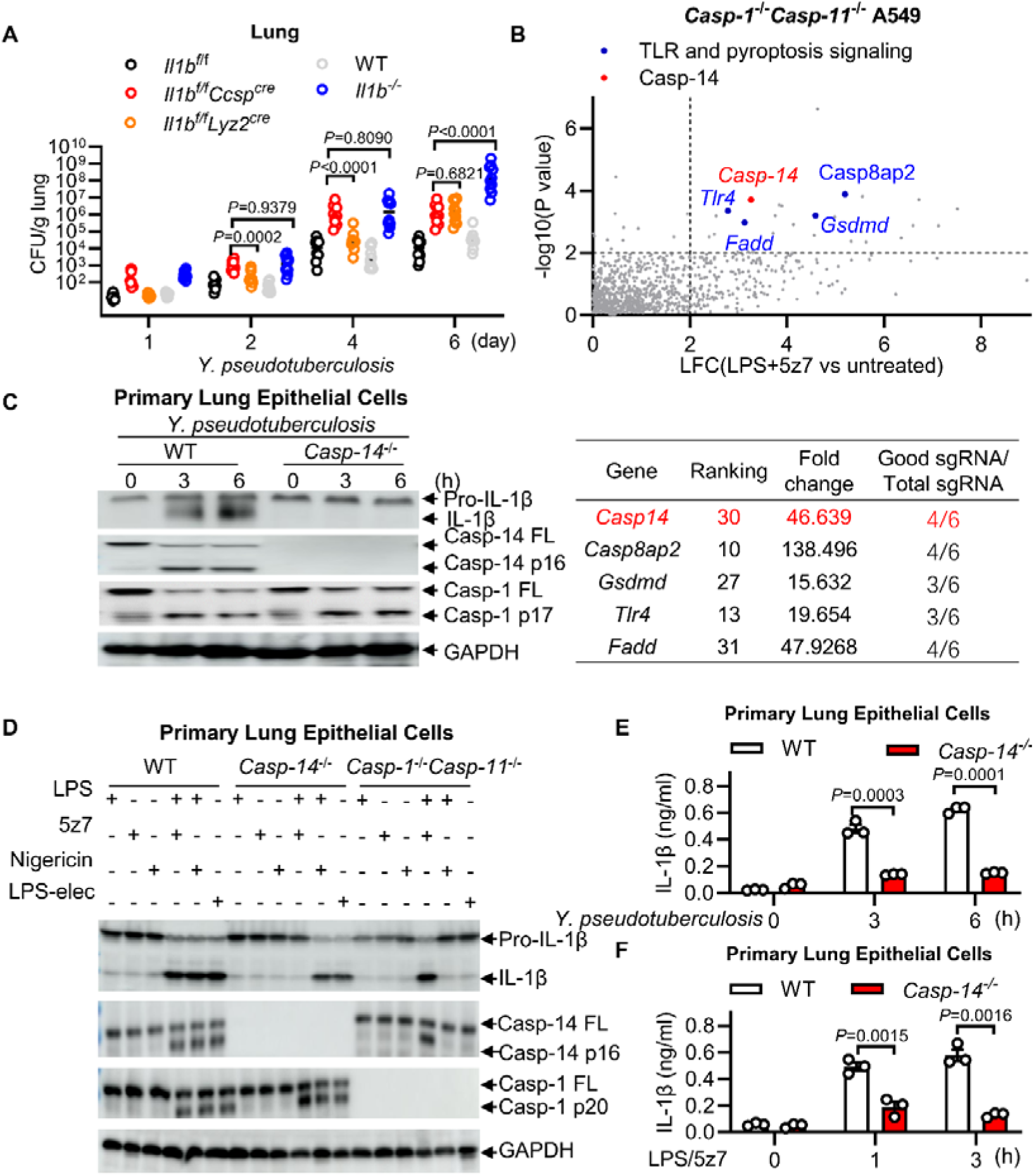
Caspase-14 processes IL-1β in Yersinia-infected epithelial cells. **(a)** 6-8 weeks female *Il1b^f/f^* mice, or lung epithelial cells conditional knock-out mice *Il1b^f/f^ Ccsp^Cre^*, or macrophages conditional knock-out mice *Il1b^f/f^ Lyz2^Cre^,* or wild-type mice, or whole-body gene knockout mice *Il1b^-/-^* aerosol challenged with *Y. pseudotuberculosis (*∼10^7^ CFU/mice). CFU in lung of at indicated days post infection. **(b)** List of sgRNA hits from the screening relative to controls in positive selection. The dotted lines represent *P* = 0.01. List of sgRNA hits from the screening in *Caspase-1*^-/-^ *Caspase-11*^-/-^A549 treated with LPS/5z-7. Corresponding genes targeted with multiple sgRNAs and fold enrichments are shown. **(c, d)** Immunoblots of WT, or *Caspase-1^-/-^Caspase-11^-/-^*, or *Caspase-14^-/-^* primary lung epithelial cells infected with *Yersinia* **(C)**, or stimulated as indicated for3 hours. **(e, f)** IL-1β release was measured by ELISA of primary lung epithelial cells from WT or *Caspase-14^-/-^*mice infected with *Y. pseudotuberculosis,* or stimulated with LPS/5z-7. All of the immunoblot data are representative images from one of three independent experiments. Results in **a**, **e** and **f** reflect the mean ± s.e.m from three independent biological experiments. Two-tailed unpaired Student’s t-test were used.

**Caspase-14 processes IL-1**β in *Y. pseudotuberculosis-*infected epithelial cells

The maturation of IL-1β is typically mediated by Caspase-1 or -11 ^11^. However, deletion of *Caspase-1* and -*11* partially attenuated the cleavage of pro-IL-1β (**Extended Data Fig.1b**), or the secretion of IL-1β (**Extended Data Fig.1c**) in lung tissues of *Y. pseudotuberculosis-*infected mice, which is consistent with previous reports ^50,51^. Consistently, *Caspase-1* and *-11* knockout led to a partial reduction of the cleavage of pro-IL-1β or secretion of IL-1β in primary lung epithelial cells stimulated with LPS plus 5z7 (**Extended Data Fig.1d-e)**, but abolished pro-IL-1β cleavage in response to canonical inflammasome activator LPS plus nigericin ^52^, or that induced by noncanonical inflammasome activator cytoplasmic LPS through LPS electroporation ^12,13^ (**Extended Data Fig.1d-e**). These results suggest that the cleavage of pro-IL-1β in *Y. pseudotuberculosis-*infected epithelial cells may go through an unknown pathway.

To search for additional mechanisms for the cleavage of pro-IL-1β during *Y. pseudotuberculosis* infection in epithelial cells, we constructed a modified transgene-encoded biosensor ^53^, in which enhanced green fluorescence protein (EGFP) was fused to the N-terminus of mouse pro–IL-1β and mCherry fused to the C-terminus (**Extended Data Fig.1f**). Cleavage and secretion of mature C-terminus IL-1β led to the attenuation of the C-terminus mCherry signal, but the EGFP signal was kept ^33^. Thus, such an IL-1β biosensor enables a fluorescence-activated cell sorting–based assay of pro-IL-1β cleavage in cells. LPS plus 5z7 stimulation led to the cleavage of pro–IL-1β along with a dramatically decreased mCherry signal in IL-1β biosensor–expressing A549 cells, a human lung epithelial cell line (**Extended Data Fig.1g**). To avoid interference by a secondary Caspase-1 signal of pro–IL-1β cleavage ^14^, *Caspase-1*^−/−^ and *-11^-/-^*A549 cells were used to perform a genome-wide CRISPR/Cas9 screen. A *Caspase-1*^−/−^ and *-11^-/-^* A549 cells–expressing pro–IL-1β biosensor was infected with lentiviruses encoding Cas9 and a library of single-guide RNAs (sgRNAs) following treatment with extracellular LPS plus 5z7. The sorted cells were gated for an EGFP and mCherry double-positive population to identify enriched sgRNAs that target genes required for the cleavage of pro-IL-1β (**Extended Data Fig.1h**). Two replicate screens were performed with different batches of *Caspase-1*^−/−^ and *-11^-/-^* A549 transduced with a gRNA library for 8 and 10 days, respectively. The screening identified *Caspase-14* as showing the top hits in the cleavage of pro-IL-1β (**Fig. 1b; Supplementary Table 1)**. In addition, none pro-IL-1β cleavage cells in LPS plus 5z7 stimulation were highly enriched for sgRNAs targeting genes involved in Toll-like receptor (TLR) or pyroptosis signaling, including *Tlr4*, *Fadd*, *Casp8ap2*, and *Gsdmd* (**Fig. 1b**). We also performed pro-IL-1β cleavage screening in WT A549 cells, as shown in **Extended Data Fig.1i**, *caspase-14* was identified among the top 50 hits involved in the cleavage of pro-IL-1β in response to LPS plus 5z7 stimulation **(Supplementary Table 2)**. Consistent with previous reports, pyroptosis or TLR signaling-related genes were also enriched including *Fadd* ^15,17^, *Caspase-8* ^15,17^, *Caspase-1*^14^, *Gsdme* ^54^, *Nlrp1b* ^55–57^ .

*Caspase-14* is reported to be highly expressed in keratinocytes or epithelial cells rather than macrophages ^8,43–45^. Indeed, our western blot analysis revealed that the protein level of Caspase-14 in BMDMs was much lower compared to epithelial cells **(Extended Data Fig.1j)**. Caspase-14 protein level was highest in A549 cells, a human lung epithelial cell line, compared to several other epithelial cell lines examined, including HeLa (human cervical epithelium) or MB49 (mouse bladder urothelium) (**Extended Data Fig.1j**). As shown in **Extended Data Fig.1k-m**, WT A549 cells, but not *CASP-14*^-/-^ A549 cells exhibited significant cleavage of pro-IL-1β or secretion of IL-1β in response to *Y. pseudotuberculosis* infection or LPS plus 5z7 stimulation. To further verify the role of caspase-14 in the *Y. pseudotuberculosis*-induced cleavage of pro-IL-1β in epithelial cells, we generated *Casp-14^em^*^1^ (*Caspase-14*^-/-^) mice that carried a global deletion of *Caspase-14* gene. *Caspase-14*^-/-^ primary lung epithelial cells showed no detectable IL-1β maturation in response to *Y. pseudotuberculosis* infection or LPS plus 5z7 stimulation (**Fig.1c, d**). Consistently, a much lower level of IL-1β was released from *Caspase-14*^-/-^ primary lung epithelial cells infected with *Y. pseudotuberculosis* (**Fig.1e**) or stimulated with LPS plus 5z7 (**Fig.1f**) as compared to those of wild-type lung epithelial cells . However, the induced cleavage of pro-IL-1β by LPS plus nigericin or LPS electroporation in lung epithelial cells was not significantly changed by the deletion of *Caspase-14* (**Fig.1d**), suggesting that cleavage of pro–IL-1β in epithelial cells infected with *Y. pseudotuberculosis* or stimulated with LPS plus 5z7 may specifically go through a Caspase-14-dependent pathway. Moreover, deletion of *Caspase-14*, but not of Caspase-1 and -11 inhibited cleavage of pro-IL-1β or secretion of IL-1β in *Y. pseudotuberculosis-*infected primary intestine epithelial cells **(Extended Data Fig.1n-q)**. These results suggest that Caspase-14-dependent cleavage of pro–IL-1β may function as a general mechanism in different epithelial cells.

### Caspase-8 processes Caspase-14 on D146

Caspases are aspartate-specific cysteine proteases whose activation and function are triggered by a delicate caspase-cascade system ^58–60^. Previous study reports that hymotrypsin-like serine protease kallikrein-related peptidase-7 (KLK7) cleaves Caspase-14 to yield the p17/p10 mature form during keratinocyte terminal differentiation ^46^. We consistently observed a cleavage product (∼16 kDa) of Caspase-14 in primary lung epithelial cells infected with *Y. pseudotuberculosis* (**Fig.2a**) or stimulated with LPS plus 5z7 (**Fig.2b**). However, no detectable cleavage of Caspase-14 was found in A549 cells stimulated with LPS plus nigericin or LPS electroporation (**Extended Data Fig.2a**), which was consistent with a nonessential role of Caspase-14 for canonical or noncanonical inflammasome–mediated cleavage of pro-IL-1β (**Fig.1d**).

**Fig. 2.**
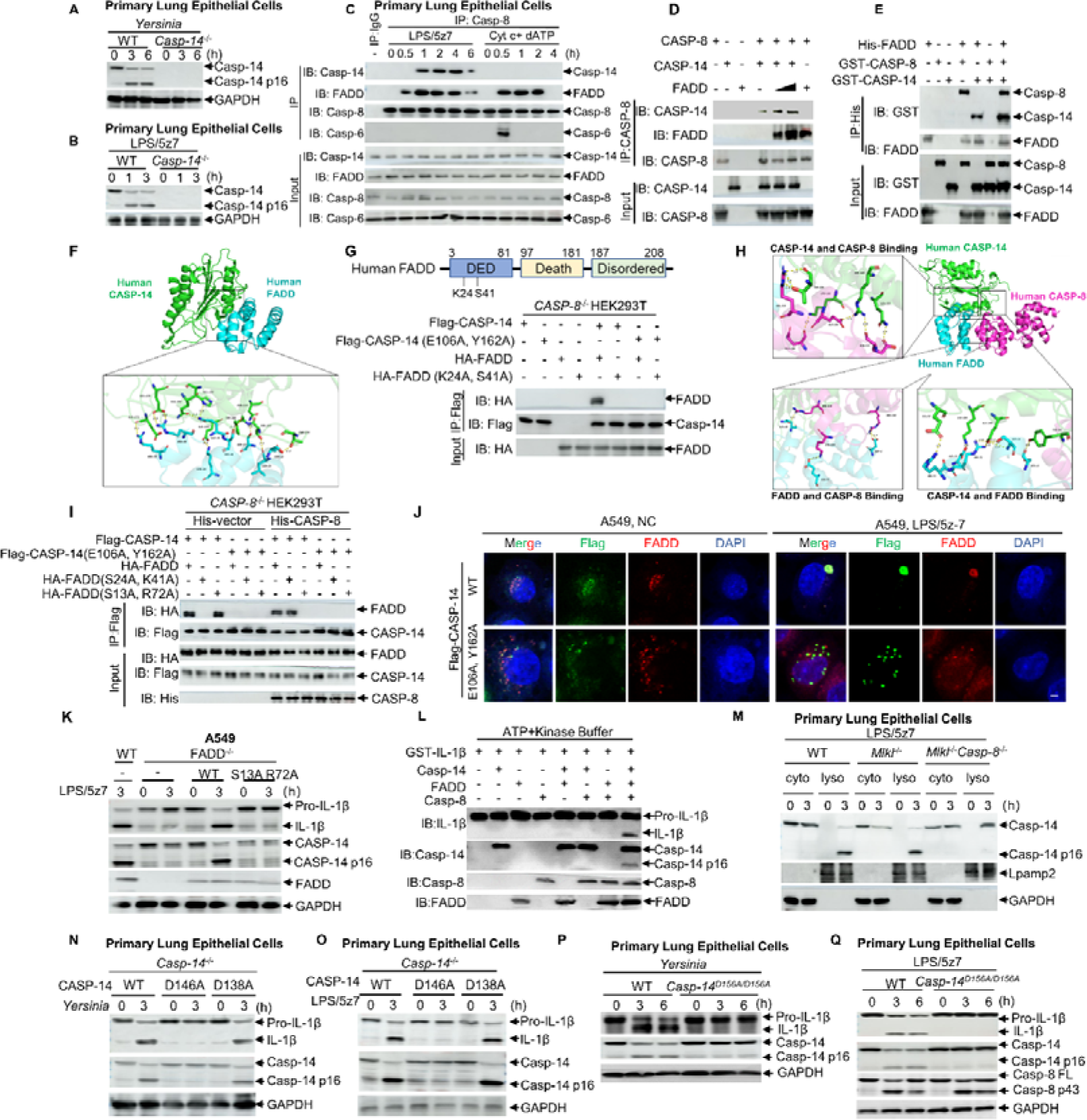
Caspase-8 processes Caspase-14 on D146. **(a, b)** Immunoblotting of indicated proteins from WT or *Caspase-14^-/-^* primary lung epithelial cells infected with *Y. pseudotuberculosis* or stimulated with LPS/5z-7 for the indicated times. **(c)** Endogenous Caspase-8 complex was immunoprecipitated with anti-Caspase-8 antibody and analyzed by immunoblot with the indicated antibodies stimulated with LPS/5z-7 or Cytochrome c + dATP for the indicated times. **(d, e**) Exogenous FADD, CASP-14 or CASP-8 was immunoprecipitated with antibody and analyzed by immunoblot with the indicated antibodies. **(f, h**) The overall structure of the complex of CASP-14 and FADD with or without CASP-8. **(g, i)** Immunoblot of the lysates and anti-Flag immunoprecipitates of *CASP-8*^-/-^ HEK293T cells stably expressed WT Flag-CASP-14 or mutants and HA-FADD or His-CASP-8. **(j)**Representative confocal fluorescence images of *CASP-14^-/-^* A549 stably expressing WT CASP-14 or mutants stimulated with LPS/5z-7 for 2h. Scale bars, 2 μm. **(k)** Immunoblotting of indicated proteins from WT or *FADD^-/-^* A549 stably expressing HA-FADD and HA-FADD (S13A and R72A) stimulated with LPS/5z-7 for the indicated times. **(l**) *In vitro* cleavage assay of pro-IL-1β protein with or without Caspase-8, Caspase-14 or FADD. **(m)** Immunoblotting of indicated proteins in lysosome fraction from WT, or *Mlkl^-/^,^-^* or *Mlkl^-/-^ Caspase-8^-/-^* primary lung epithelial cells stimulated with LPS/5z-7 for the indicated times. **(n, o)** Immunoblotting of indicated proteins from WT or *Caspase-14^-/-^* primary lung epithelial cells expressing WT Caspase-14 or mutants, infected with *Y. pseudotuberculosis* or stimulated with LPS/5z-7 for the indicated times. **(p, o**) Immunoblotting of indicated proteins from WT or *Caspase-14*^D156A/D156A^ primary lung epithelial cells infected with *Y. pseudotuberculosis,* or stimulated with LPS/5z-7. All of the immunoblot data are representative images from one of three independent experiments. Two-tailed unpaired Student’s t-test were used.

We next investigated the molecular mechanism underlying the cleavage of caspase-14. By performing co-immunoprecipitation, we found that Caspase-14 interacted with FADD and Caspase-8 in LPS plus 5z7–stimulated primary lung epithelial cells or A549 cells (**Fig.2c and Extended Data Fig.2b**). The presence of FADD protein increased the binding affinity of Caspase-8 and Caspase-14 *in vitro* (**Fig.2d**). It has been shown that Caspase-8 binds to FADD through a classical DED-DED interaction, which caused oligomerization and activation of Caspase-8 ^61,62^. However, Caspase-14 does not possess a DED domain as Caspase-8 does ^8,41^, but showed similar affinity with FADD as Caspase-8 protein did *in vitro* (**Fig.2e**). To further improve the logical consistency of Caspase-8/FADD/Caspase-14 interaction, we utilized the specialized protein-protein docking program HDOCK for docking with human CASP-14 (uniprot ID: P31944) and Human FADD (uniprot ID: Q13158). Multiple hydrogen bonds were found to be formed between the human CASP-14 protein and human FADD protein, in which E106 & Y162 of human CASP-14 and K24 & S41 of human FADD protein were involved (**Fig. 2f**). To investigate the interaction, we performed immunoprecipitation (IP) assays in HEK293T cells expressing Flag-tagged WT CASP-14 or the E106A & Y162A mutant. These cells were co-transfected with or without HA-tagged WT FADD or the K24A & S41A mutant. We found that mutations at the CASP-14 binding sites (E106A & Y162A) or the FADD binding sites (K24A & S41A) impaired the CASP-14–FADD interaction (**Fig. 2g**). Therefore, these findings suggest that though Caspase-14 does not have a DED domain, E106 & Y162 on Caspase-14 may interact with K24 & S41 on the DED domain of FADD.

We further analyzed the spatial interaction forms among three proteins: Human CASP-14 (uniprot ID: P31944), Human FADD (uniprot ID: Q13158) and Human CASP-8 (uniprot ID: Q14790). At the presence of CASP-8, while FADD-interacting amino acids of CASP-14 remained to be E106 & Y162 constantly (**Fig.2h-j**), but the CASP-14-interacting sites of FADD switched from K24 & S41 to S13 & R72 (**Fig. 2h-j**), which were essential for the cleavage of Caspase-14 in LPS plus 5z7-stimulated A549 cells (**Fig.2k**). Consistent with previous report ^63^, when Caspase-14 was present, FADD still interacted with Caspase-8 through D2 & E51 on the DED domain (Fig. 2h). These results suggest that different sites of FADD may interact with Caspase-14 and Caspase-8 in the complex, thus leading to the cleavage of Caspase-14.

Given that Caspase-14 interacts with Caspase-8 and FADD, we next examined whether Caspase-8 could regulate the cleavage of Caspase-14. Purified recombinant Caspase-14 (∼28 kDa) was incubated with Caspase-8 and FADD protein, and Caspase-14 was found to be cleaved, forming a ∼16 kDa C-terminal fragment (p16) (**Fig. 2l**). Consistently, no detectable cleavage of Caspase-14 was detected in *Caspase-8-* (**Extended Data Fig.2c**) or *Fadd-* (**Extended Data Fig.2d**) deficient primary lung epithelial cells stimulated with LPS plus 5z7. However, in LPS plus 5z7-stimulated primary lung epithelial cells (**Extended Data Fig.2e**), deletion of *Gsdmd* did no effect on Caspase-14 cleavage indicating that cleavage of Caspase-14 may not relate to GSDMD-mediated pyroptosis. During *Y. pseudotuberculosis* infection, Caspase-8 was recruited and activated on lysosomes ^16^. Cleavage of caspase-14 was detected in the lysosome protein fraction from primary lung epithelial cells stimulated with LPS plus 5z7, but deletion of Caspase-8 abolished the cleavage of Caspase-14 on lysosomes (**Fig. 2m**). These results suggest that activated caspase-8 may cleave caspase-14 on lysosomes in response to *Y. pseudotuberculosis* infection.

Sequence analysis of substrate recognition motifs of Caspase-8 ^64^ suggests that Caspase-8 may cleave Caspase-14 after Asp at 146, within _141_VGGD_146_ in human Caspase-14, or _153_LGGD_156_ in mouse Caspase-14 (**Extended Data Fig.2f**). We thus generated a mutant Caspase-14, in which D146 (D156 in mouse Caspase-14) was replaced with an alanine. Indeed, CASP-14^D146A^ was resistant to cleavage by CASP-8 *in vitro* (**Extended Data Fig.2g**). Consistently, *Caspase-14*^-/-^ primary lung epithelial cells expressing Caspase-14 (D146A) mutant did not show the cleavage of pro-IL-1β or the secretion of IL-1β as those expressing WT Caspase-14 did in response to *Y. pseudotuberculosis* infection or LPS plus 5z7 stimulation (**Fig.2n, o and Extended Data Fig.2h, i**). Furthermore, we generated *Casp-14^D156A^* knock-in mice. *Casp-14*^D156A^ primary lung epithelial cells showed no detectable cleavage of Caspase-14 in response to *Y. pseudotuberculosis* infection or LPS plus 5z7 stimulation (**Fig.2p, q**). In contrast to WT cells, *Casp-14*^D156A^ mutant lung epithelial cells did not show the cleavage of pro-IL-1β or release IL-1β upon either *Y. pseudotuberculosis* infection or treatment with LPS and 5z7 (**Fig.2p, q and Extended Data Fig.2j, k**). Those findings suggest that caspase-8 may specifically cleave caspase-14 at D146/156 within a _141_VGGD_146_ (or _153_LGGD_156_) motif.

### Processing of pro–IL-1β by Caspase-14 p16

It has been shown that a high concentration of active fragment of Caspase-8 (300ng) can directly cleave pro-IL-1β *in vitro* ^65^. Other studies and our data showed the deficiency of pro-IL-1β cleavage in *Caspase-8* knockout cells ^26–28^ **(Extended Data Fig.2C**). To further define the function role of Caspase-8 and Caspase-14 on pro–IL-1β processing, the cleavage of pro–IL-1β was examined with increasing dose (30ng, 100ng and 300ng) of full length of Caspase-8 or Caspase-14 plus FADD protein *in vitro* (**Extended Data Fig.3a**). The FADD protein triggers autoproteolytic activation of full-length Caspase-8 (30 ng) to generate the active fragment Caspase-8 p43 *in vitro* ^66^, which is sufficient to cleave Caspase-14 into active Caspase-14-p16 (**Extended Data Fig.3a**), but insufficient to cleave pro-IL-1β *in vitro* (**Extended Data Fig.3a**). Much higher level of full-length Caspase-8 (300ng) plus FADD was required for cleavage of pro–IL-1β *in vitro* (**Extended Data Fig.3a**). In contrast, full-length Caspase-14 (30 ng), when cleaved by Caspase-8 p43 (from 30ng full-length caspase-8) to generate active Caspase-14 p16, was sufficient to cleave pro-IL-1β *in vitro* (**Extended Data Fig.3a**). Furthermore, active Caspase-14-p16 protein (30ng), a concentration similar to the commonly used concentration of active Caspase-1 p20 for the cleavage of pro–IL-1β ^67^, plus FADD or kosmotropic salts ^7^, was sufficient to cleave pro–IL-1β *in vitro* (**Extended Data Fig.3b**). These results suggest that Caspase-14, not Caspase-8, may act as a predominant processor of pro–IL-1β.

Other studies have shown the deficiency of IL-1β cleavage in *Caspase-8* knockout cells ^26–28^. Indeed, *Caspase-8* knockout diminished the cleavage of pro-IL-1β in primary lung epithelial cells stimulated with LPS plus 5z7 (**Extended Data Fig.2c**). However, although Caspase-8 was cleaved and active in *Casp-14*^D156A^ knock-in primary lung epithelial cells (a cleavage-resistant mutant), pro–IL-1β processing was not detectable following *Y. pseudotuberculosis* infection. (**Fig.2p, q**). These data suggested that Caspase-8 may mediate the cleavage of pro–IL-1β more efficiently through Caspase-14. Furthermore, *Y. pseudotuberculosis* infection or LPS plus 5z7 stimulation induced the cleavage of pro–IL-1β in *Caspase-14*^-/-^ primary lung epithelial cells complemented with WT Casp-14, but not Casp-14^D146A^ mutant, which is unresponsive to Caspase-8-dependent cleavage (**Fig.2n, o**). Consistently, we did not detect secretion of mature IL-1β protein in the supernatant of primary lung epithelial cells expressing Casp-14*^D146A^* or Casp-14*^D156A^* mutant in response to *Y. pseudotuberculosis* infection or LPS plus 5z7 stimulation (**Extended Data Fig.2g-j**). Together, these findings suggest that Caspase-8 may actually cleave Caspase-14 into an active Caspase-14 p16 fragment, which subsequently processes pro-IL-1β under physiological conditions.

Although protein-protein interaction databases suggested that human Caspase-8 could also interact with Caspase-1/2/3/4/6/7/9/10/14 (https://thebiogrid.org/107291), we only detected the interaction of Caspase-1/3/14 with Caspase-8 in A549 cells stimulated with LPS plus 5z7 (**Extended Data Fig.3c**). Furthermore, we examined cleavage of pro-IL-1β *in vitro* by active Caspase-1/2/3/4/6/7/8/9/10/14 (30ng individually) fragment plus FADD protein. Only active Caspase-1 p20 fragment or active Caspase-14 p16 fragment showed significant processing of pro-IL-1β protein *in vitro* (**Extended Data Fig.3d**). In LPS plus 5z7 stimulated A549 cells, knockdown of Caspase-3 by specific siRNA or deletion of *Caspase-1* had no significant effects on the cleavage of pro-IL-1β (**Extended Data Fig.3e, f**). However, *Caspase-14* knockout nearly completely blocked the IL-1β secretion at 1h and 3h after LPS plus 5z7 stimulation in primary lung epithelial cells (**Extended Data Fig.3f**), while maintaining cell viability comparable to control cells (**Extended Data Fig.3g**). These results suggest that Caspase-14 in the death complex may be responsible for the cleavage of pro-IL-1β in response to LPS plus 5z7 stimulation in epithelial cells.

Given that Caspase-14 is mainly responsible for the cleavage of pro-IL-1β, we next investigated how the Caspase-14 p16 fragment executes IL-1β processing. Caspase-14 immunoprecipitated from LPS plus 5z7–stimulated primary lung epithelial cells, but not from LPS plus nigericin–stimulated or LPS-electroporated macrophages, was sufficient for the cleavage of purified recombinant pro–IL-1β *in vitro* (**Fig.3a**). In addition, purified Caspase-14 p16 from LPS plus 5z7–stimulated epithelial cells directly cleaved recombinant pro–IL-1β into a mature IL-1β p17 fragment *in vitro* (**Fig.3b**). However, prokaryotic purified human active CASP-14 p16 fragment exhibited much higher affinity with pro–IL-1β protein than prokaryotic purified human active CASP-1 p20 fragment (**Fig.3c**). Previous positional scanning substrate library analysis revealed the preferences substrate of human Caspase-14 matches the specificity of the cytokine activator Caspases-1, -4, and -5, while the mouse enzyme shows more similarity toward apical Caspases-8 and -9 ^7,68,69^. However, the natural substrate of human or murine Caspase-14 remains elusive ^7^. To improve the logical consistency of different substrate preferences between human caspase-14 and murine Caspase-14, we examined the cleavage of pro-IL-1β by the prokaryotic purified active fragment p16 of human or murine Caspase-14 *in vitro,* and found that human or murine Caspase-14 p16 fragment had similar cleavage activity towards pro-IL-1β in the buffer supplemented with kosmotropic salts (**Extended Data Fig.3H**) ^7^. Consistently, we also observed the cleavage of pro-IL-1β in *CASP-14*^−/−^ A549 cells stably expressing human or murine Caspase-14 at 3h post LPS plus 5z7 stimulation (**Extended Data Fig.3i**). These results suggest that both human and murine active Caspase-14 p16 fragment may directly cleave pro–IL-1β.

**Fig. 3.**
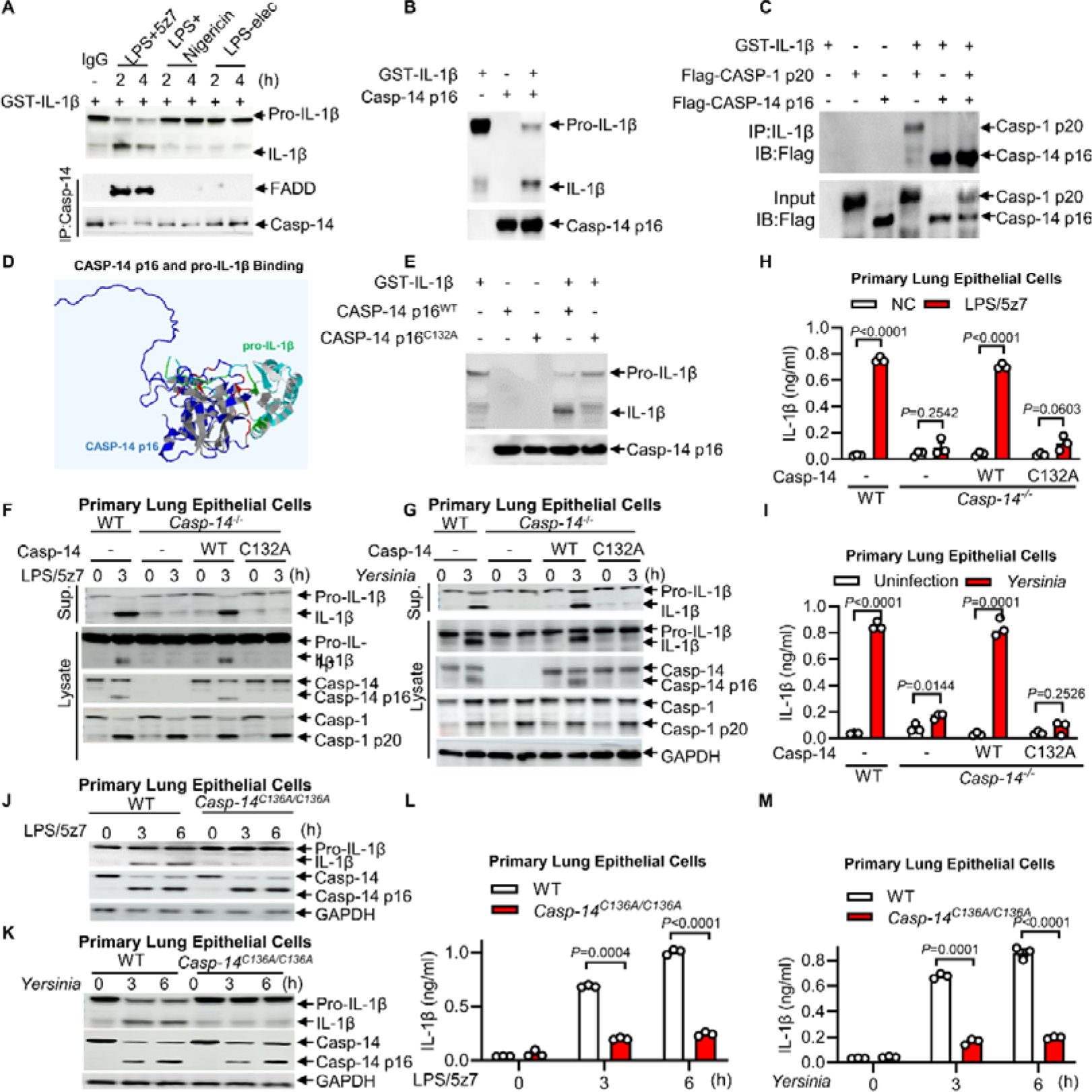
Processing of pro–IL-1β by Caspase-14 p16. **(a)** *In vitro* cleavage assay of pro-IL-1β protein with caspase-14 isolated from primary lung epithelial cells stimulated with LPS electroporation, LPS/nigericin or LPS/5z-7 for the indicated times. **(b, e**) *In vitro* cleavage assay of pro-IL-1β protein with or without Caspase-14 p16 fragment, Caspase-14 p16 ^C132A^ fragment. **(c**) Exogenous Caspase-14 p16 fragment or Caspase-1 p17 fragment isolated from HEK293T cells was immunoprecipitated with anti-Flag antibody and analyzed by immunoblot with the indicated antibodies. **(d**) The overall structure of the complex of CASP-14 p16 and pro-IL-1β. **(f, g**) Immunoblotting of indicated proteins from WT or *Caspase-14^-/-^* primary lung epithelial cells expressing WT Caspase-14 or mutant, and infected with *Y. pseudotuberculosis* or stimulated with LPS/5z-7 for the indicated times. **(h, i)** IL-1β release was measured by ELISA from WT or *Caspase-14^-/-^* primary lung epithelial cells expressing WT Caspase-14 or mutant infected with *Y. pseudotuberculosis,* or stimulated with LPS/5z-7. **(j, k**) immunoblotting of indicated proteins from WT or *Casp-14^C136A/C136A^* primary lung epithelial cells infected with *Y. pseudotuberculosis* or stimulated with LPS/5z-7 for the indicated times. **(l, m)** IL-1β release was measured by ELISA from WT or *Casp-14^C136A/C136A^*primary lung epithelial cells infected with *Y. pseudotuberculosis* or stimulated with LPS/5z-7 for the indicated times. All of the immunoblot data are representative images from one of three independent experiments. Results in **h**, **i, l** and **m** reflect the mean ± s.e.m from three independent biological experiments. Two-tailed unpaired Student’s t-test were used.

Structural simulation of molecular docking between human CASP-14 p16 and pro–IL-1β protein revealed that Cys132 in the CASP-14 p16 fragment formed a salt bridge with the cleavage region of pro–IL-1β (**Fig.3d**). Notably, Cys132 is highly conserved between mouse or human Caspase-14 **(Extended Data Fig.2f)**, suggesting that this enzyme active site may determine the protease activity of Caspase-14. Thus, we generated a human CASP-14 p16 fragment mutant in which Cys132 (Cys136 in mouse Caspase-14 p16 fragment) was replaced with an alanine. C132A mutants of human CASP-14 p16 fragment was unable to cleave recombinant pro–IL-1β *in vitro* (**Fig.3e**). *Caspase-14*^-/-^ A549 cells and primary lung epithelial cells complemented with a human CASP-14 (C132A) mutant exhibited no cleavage of pro–IL-1β and mature IL-1β secretion in response to *Y. pseudotuberculosis* infection or LPS plus 5z7 stimulation (**Fig.3f, g and Extended Data Fig.3j, k**). Consistent with this, no significant release of mature IL-1β was detected in the supernatant of cultured A549 cells or primary lung epithelial cells expressing human CASP-14 (C132A) mutant in response to *Y. pseudotuberculosis* infection or LPS plus 5z7 stimulation (**Fig.3h, i and Extended Data Fig.3l, m**). We further constructed *Casp-14*^C136A^ knock-in mice. Moreover, no cleavage of pro–IL-1β or mature IL-1β secretion were detected in primary lung epithelial cells from *Casp-14*^C136A^ mice in response to *Y. pseudotuberculosis* infection or LPS plus 5z7 stimulation (**Fig.3j-m**). These data indicated Cys132/136 in Caspase-14 p16 as a protease active site.

### TAK1 phosphorylates Caspase-1 at S126

Phylogenetic tree analysis traces Caspase-14, but not Caspase-1, back to the anole lizard and *Anolis carolinensis*, which indicates that Caspase-14 is more evolutionarily conserved than Caspase-1 ^70^. Given that both Caspases-14 and -1 active fragments can cleave pro-IL-1β, we next investigated how they are differentially regulated. It has been shown that TAK1 is a serine/threonine kinase ^71,72^, and *Y. pseudotuberculosis* YopJ or the small molecule compound 5z7 inhibits TAK1’s kinase activity ^14,15,21,22,25,37^. *Caspase-1* deletion abolished pro-IL-1β cleavage induced by LPS plus nigericin in primary lung epithelial cells (**Extended Data Fig.4a**). Conversely, TAK1 inhibition with 5z7 restored pro-IL-1β cleavage and induced Caspase-14 p16 in *Caspase-1*^-/-^*Caspase-11*^-/-^ epithelial cells (**Extended Data Fig.4a**). While *Caspase-14* deletion did not affect LPS plus nigericin induced pro-IL-1β cleavage, it blocked pro-IL-1β cleavage in cells treated with LPS plus nigericin and 5z7 (**Extended Data Fig.4b**). Interestingly, we observed the cleavage of Caspase-1 into an active Caspase-1 p20 form, but not the cleavage of pro–IL-1β in *Caspasae-14*^-/-^ A549 cells or epithelial cells infected with *Y. pseudotuberculosis* or stimulated with LPS plus 5z7 (**Fig.1c and Extended Data Fig.1k**), suggesting that active Caspase-1 p20 may need to be activated by TAK1 first to execute the cleavage of pro-IL-1β. Indeed, addition of TAK1 plus TAB1 (TAK1-TAB1), which induced kinase activity of TAK1 ^73,74^, resulted in the phosphorylation of Caspase-1 *in vitro* as detected by western blot analysis using a pan-phosphoserine/threonine antibody (**Fig.4a**). Through mass spectrometry analysis we identified two phosphorylation sites on Caspase-1 p20 fragment by TAK1 *in vitro*, including S126 and T230. However, only Caspase-1 (S126A) mutant showed no detectable phosphorylation by the pan-phosphoserine/threonine antibody when incubated with TAK1 plus TAB1 *in vitro* (**Fig.4a**), suggesting that TAK1 may mainly phosphorylate Caspase-1 p20 fragment on Ser 126. We further generated a Caspase-1 S126 phosphorylation-specific antibody by immunizing a rabbit with c(KLH)-PT(S-p)-SGSEGN peptide. *In vitro* experiments showed that Caspase-1 S126 phosphorylation was induced by the TAK1–TAB1 complex, but not by TAK1 or TAB1 alone (**Fig.4b**). In addition, *Caspase-1*^/^ A549 cells stably expressing Caspase-1 (S126A) mutant, but not WT Caspase-1 or Caspase-1 (T230A) mutant exhibited normal autocleavage of Caspase-1, but showed no detectable cleavage of pro-IL-1β upon stimulation with LPS plus nigericin—a canonical Caspase-1 activator (**Fig.4c**). These results suggest that Caspase-1 activation and subsequent pro-IL-1β cleavage may require TAK1-mediated phosphorylation of Caspase-1 at S126. Consequently, blockade of TAK1 by the *Y. pseudotuberculosis* effector YopJ likely inhibits Caspase-1-dependent pro-IL-1β cleavage, thereby redirecting processing to a Caspase-14-dependent pathway.

**Fig. 4.**
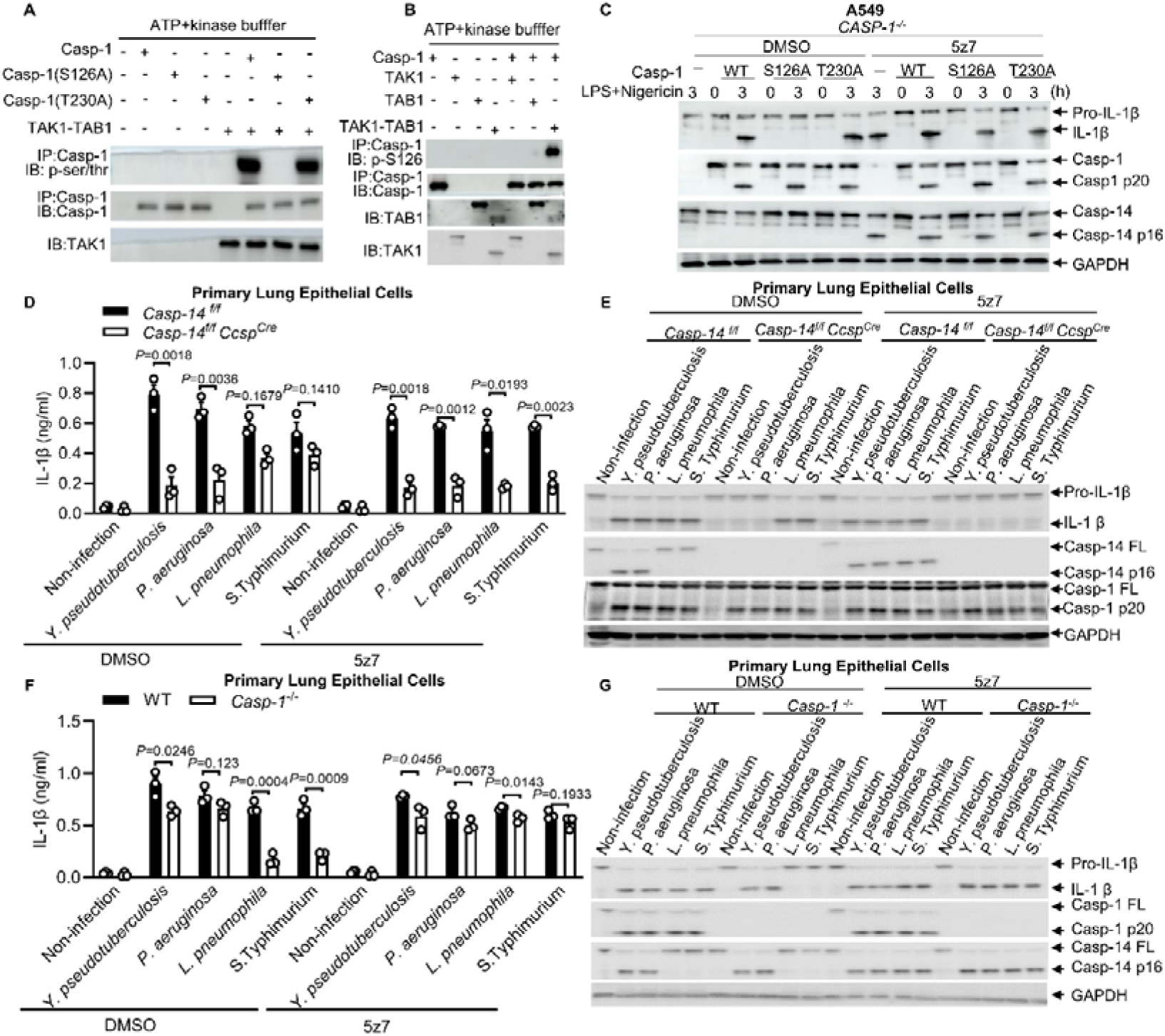
TAK1 phosphorylates Caspase-1 at S126. **(a, b)** *In vitro* kinase assay of purified recombinant GST-TAK1, TAB1 or TAK1-TAB1 (active) with WT Caspase-1 or mutants. **(b)** Immunoblotting of indicated proteins from *CASP-1^-/-^* A549 stably expressing WT Caspase-1 or mutants, and stimulated with LPS/nigericin for the indicated times in the absence or presence of 5z-7. **(d, f**) IL-1β release was measured by ELISA from primary lung epithelial cells infected with *Y. pseudotuberculosis, P. aeruginosa L. pneumophila,* or *S. Typhimurium* (MOI=5) for 12h. **(e, g**) Immunoblotting of indicated proteins from primary lung epithelial cells infected with *Y. pseudotuberculosis, P.aeruginosa*, *L. pneumophila,* or *S. Typhimurium* (MOI=5) for 12h. All of the immunoblot data are representative images from one of three independent experiments. Results in **D** AND **F** reflect the mean ± s.e.m from three independent biological experiments. Two-tailed unpaired Student’s t-test were used.

We next examined the contribution of Caspase-1 or Caspase-14 to the cleavage of pro–IL-1β during other pathogenic bacterial infections. In primary lung epithelial cells, Caspase-14 was essential for pro-IL-1β cleavage and IL-1β secretion in response to *P. aeruginosa* or *Y. pseudotuberculosis* infection, but not *L. pneumophila, S. Typhimurium* infection, suggesting Caspase-14-mediated cleavage of pro–IL-1β as a general mechanism for different infectious diseases (**Fig.4d, e**). Interestingly, our western blot analysis revealed that phosphorylation of TAK1 was also blocked by *P. aeruginosa* infection as did by *Y. pseudotuberculosis* infection **(Extended Data Fig.4c)** ^75^, indicating a correlation between TAK1 blockade and Caspase-14-dependent cleavage of pro–IL-1β during *P. aeruginosa* or *Y. pseudotuberculosis* infection. Moreover, inhibition of TAK1 by 5z7 abolished the cleavage of pro–IL-1β possibly by Caspase-1 in *Caspase-14^-/-^* epithelial cells infected with *L. pneumophila* or *S. Typhimurium* infection (**Fig.4d, e**), but led to the cleavage of pro-IL-1β by Caspase-14 in *Caspase-1*^-/-^ primary lung epithelial cells (**Fig.4f, g**). Together, these results suggest that while phosphorylation of Caspase-1 by TAK1 may trigger Caspase-1-dependent cleavage of pro–IL-1β in response to *L. pneumophila* or *S. Typhimurium* infection, blockade of TAK1 by bacterial effector may lead to the Caspase-14-dependent cleavage of pro-IL-1β in response to *P. aeruginosa* or *Y. pseudotuberculosis* infection. Thus, our findings revealed different cleavage of pro-IL-1β modes mediated by the molecular switch of TAK1 activation in response to different pathogenic bacterial infection.

### Caspase-14-dependent maturation of IL-1β determines anti-bacterial immunity

To verify the contribution of Caspase-14 to the *Y. pseudotuberculosis*-induced cleavage of pro-IL-1β, we crossed *Caspase-14^flox/flox^* (*Casp-14^f/f^*) mice with Ccsp-Cre transgenic mice and generated *Caspase-14*^flox/flox^; Ccsp-Cre (*Casp-14^f/f^ Ccsp^Cre^*) mice that carried a specific deletion of *Caspase-14* gene in lung epithelial cells ^49^. Upon aerosol challenged with *Y. pseudotuberculosis*, *Casp-14^f/f^ Ccsp^Cre^* mice exhibited dramatically decreased IL-1β production in their serum (**Fig.5a**) and lung tissues (**Fig.5b**), indicating that epithelial Caspase 14 contributes to IL 1β activation during infection. Aerosol challenge with *Y. pseudotuberculosis* resulted in a significantly higher bacterial load in the lungs of *Il1b*^/^ mice (**Fig.5c**), demonstrating the essential role of IL-1β in controlling *Y. pseudotuberculosis* in the lungs. Consistent with these findings, *Casp-14^f/f^ Ccsp^Cre^* mice showed dramatically increased lung bacterial colony-forming units and histological damage in their lungs at 3 days post-infection as compared to those control mice (**Fig.5c, d**). However, administration of an anti IL 1β neutralizing antibody to *Y. pseudotuberculosis* infected *Casp-14^f/f^ Ccsp^Cre^*mice led to a rescue of the impaired bacterial control associated with Caspase 14 deletion in lung epithelial cells (**Fig.5c**). Moreover, we found that Caspase-14 C132A (enzyme inactive mutant) knock-in mice had dramatically increased lung bacterial colony-forming units and histological damage in lungs, along with reduced IL-1β production, at 3 days post *Y. pseudotuberculosis* infection (**Fig.5d,f, g**). Specific loss of Caspase-14 in intestinal epithelial cells (*Casp-14^f/f^ Villin^cre^*) was achieved by crossing *Casp-14^f/f^* mice with *Vil1*^cre^ (also known as Villincre) mice ^76^. We found that the mice carrying a specific deletion of *Caspase-14* in intestinal epithelial cells as well as Caspase-14 C136A knock-in had dramatically increased intestine bacterial colony-forming units and histological damage in the intestine at 3 days post-oral infection **(Extended Data Fig.5a-e)**. These results suggest that Caspase-14-dependent maturation of IL-1β in epithelial cells may contribute to the inflammation and host defense against *Y. pseudotuberculosis* infection.

**Fig. 5.**
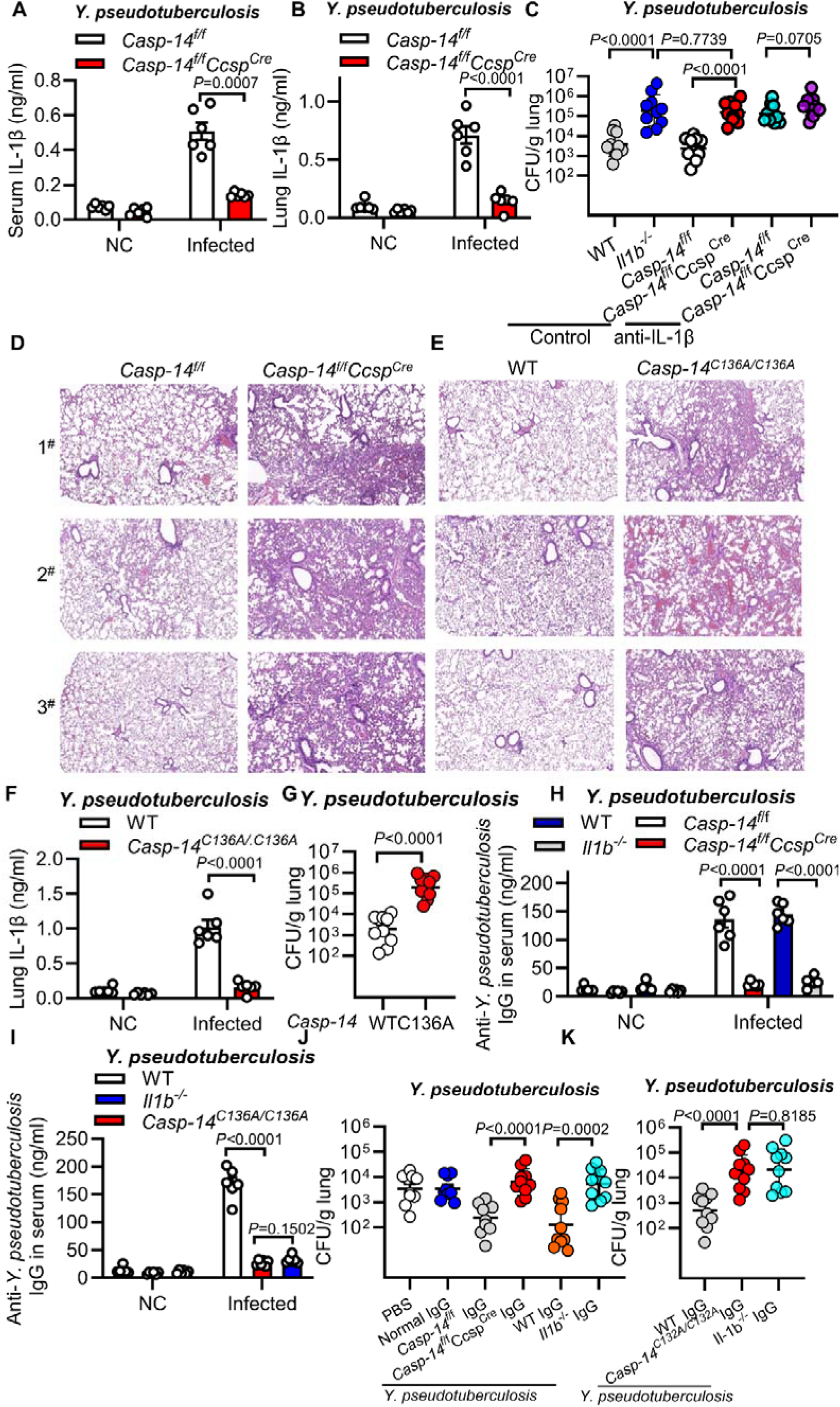
Caspase-14-dependent maturation of IL-1β protects host against infection. **(a-c)** 6-8 weeks female *Casp-14^f/f^* or *Casp-14^f/f^ Ccsp^Cre^* mice aerosol challenged with 1. *Y. pseudotuberculosis* (∼10^7^ CFU/mice). IL-1β in serum (**a**) or Lung (**b**), and CFU in lung (**C**) were detected at 3 days post infection. **(d)** H&E staining of lung tissues from *Casp-14^f/f^*, or *Casp-14^f/f^ Ccsp^Cre^* mice, or WT, or *Casp-14^C132A/C132A^*mice aerosol challenged with *Y. pseudotuberculosis* for 3 days. (scale bar, 200 μm) **(e)** 6-8 weeks female *Casp-14^f/f^* or *Casp-14^f/f^ Ccsp^Cre^*mice aerosol challenged with *Y. pseudotuberculosis* (∼10^7^ CFU/mice) with or without anti-IL-1β. CFU in lung were detected at 3 days post infection. **(f-g)** 6-8 weeks female WT or *Casp-14^C132A/C132A^*mice aerosol challenged with *Y. pseudotuberculosis* (∼10^7^ CFU/mice). IL-1β (**f**) and CFU in lung **(g)** were detected at 3 days post infection. **(h-i)** Indicated mice were immunized with *Y. pseudotuberculosis* (∼10^7^ CFU/mice). The serum concentration of IgG in the indicated mice. Data are representative of at least three independent experiments. **(j-k)** C57BL/6J mice were tail vein injected by normal mouse IgG or IgG isolated from *Y. pseudotuberculosis* infected WT or conditional knock-out Caspase-14 in lung epithelial cells or knock-in Caspase-14C136A mice (*Y. pseudotuberculosis* WT/*Il-1b^-/-^/Casp-14 KO/ KI* IgG) (15 mg/kg mouse) for 24 hours followed by aerosol injected with 1× 10^7^ CFU of *Y. pseudotuberculosis.* Results in **a, b, f, h** and **i** reflect the mean ± s.e.m from three independent biological experiments. Data were analyzed using a two-tailed Student’s t test.

Moreover, we also aerosol-infected *Casp-14^f/f^*and *Casp-14^f/f^ Ccsp^Cre^* mice with *P. aeruginosa*. As shown in **Extended Data Fig.5f, g**, *Casp-14^f/f^ Ccsp^Cre^* mice that had specific deletion of *Caspase-14* in lung epithelial cells were more susceptible to *P. aeruginosa* infection, with an uncontrolled bacterial burden in lung tissues at 5 days post infection, with dramatically lower IL-1β production. As caspase-14 is highly expressed in keratinocytes and skin tissue ^8,43–45,77^, we further examine the functional role of Caspase-14 in *Candida albicans* (*C. albicans*) skin infection model utilizing Caspase-14 fibroblast cells conditional knockout mice (*Casp-14^f/f^ Col1a2^Cre^*) ^78^. As shown in **Extended Data Fig.5h, i**, the skin tissue from *Casp-14^f/f^ Col1a2^Cre^* mice showed a much higher number of CFU and IL-1β compared those counterparts from *Casp-14^f/f^* mice. These results suggest that Caspase-14–mediated processing of IL-1β in epithelial cells may generally determine anti-bacterial immunity *in vivo*.

Humoral responses are regulated through an IL-1–dependent axis involving follicular T cell control of B cell function ^79–81^. *Y. pseudotuberculosis* induced a higher level of anti-*Y. pseudotuberculosis* IgG in wild-type mice (**Fig.5h, i**). However, deletion of IL-1β almost totally blocked the antibody production from *Y. pseudotuberculosis*-infected mice (**Fig.5h, i**), which mirrored the phenotype observed in *Il1b*^-/-^ mice (**Fig.5c**). Moreover, conditional knock-out *Caspase-14* in lung epithelial cells or knock-in *Caspase-14^C136A^*almost eliminated the total antibody production (**Fig.5h, i**) and conferred less protection against *Y. pseudotuberculosis* infection in mice (**Fig.5c, g**), which was consistent with the much lower level of IL-1β observed in those conditional *Caspase-14* knock-out or *Caspase-14^C136A^* mutant knock-in mice infected with *Y. pseudotuberculosis* (**Fig.5a, b, f**). To evaluate the effects of anti-*Y. pseudotuberculosis* IgG on *Y. pseudotuberculosis* infection, *Y. pseudotuberculosis*-infected wild-type C57BL/6J mice were administered with IgG isolated from different types of mice (**Extended Data Fig.5j**). We found that IgG from *Il1b*-deficient mice, even after prior *Y. pseudotuberculosis* exposure, failed to confer protection against infection (**Fig. 5j**), indicating that IL-1β is essential for the generation of protective anti-*Y. pseudotuberculosis* antibodies. Consistently, recipient mice transferred with IgG from lung epithelial-specific deletion of *Caspase-14* or from *Caspase-14*^C136A^ knock-in mice exhibited significantly higher bacterial loads post-infection compared to those receiving IgG from wild-type mice (**Fig. 5j, k**). These results imply that epithelial Caspase-14-dependent IL-1β maturation is a key determinant of protective humoral immunity against *Y. pseudotuberculosis*, a finding with potential relevance for vaccine development, albeit within the constraints of the mouse model used.

## Discussion

Caspases-mediated processing of cytokines coordinates cell autonomous defenses and induction of systemic inflammation^2,3,10^. Caspase-14 is typically specialized in epidermal differentiation^7,8^. Here, we establish caspase-14 as a direct processor for IL-1β maturation downstream of caspase-8–mediated pyroptosis, highlighting the critical importance of caspase-14 in controlling inflammatory responses. In contrast to canonical caspase-1-mediated IL-1β maturation that occurs mainly in monocytes ^2,3,10^, caspase-14-mediated IL-1β maturation operates more prevalently in epithelial cells.

Our data demonstrated that caspase-8–dependent cleavage of caspase-14 at Asp 146 generated a 16-kDa active fragment, which exposed the catalytic pocket necessary for cleavage of the same tetrapeptide site on pro–IL-1β as caspase-1 did, leading to IL-1β maturation. As shown in **Extended Data Fig. 6**, a surprising discovery is that the phosphorylation of S126 on caspase-1 by TAK1 is a prerequisite for its activation and subsequent cleavage of pro-IL-1β, blockade of TAK1 by *Y. pseudotuberculosis* effector YopJ inhibits caspase-1-mediated cleavage of pro-IL-1β, and attenuates TAK1/inhibitor of nuclear factor-κΒ kinase (IKK)-induced expression of cellular FLICE-like inhibitory protein (cFLIP) to restore the activation of caspase-8 ^17^, thus switching to caspase-14-dependent cleavage of pro-IL-1β. Interestingly, caspase-14 was also essential for the cleavage of pro-IL-1β and secretion of IL-1β in epithelial cells, which is essential for host inflammation and immunity against *P. aeruginosa* and *C. albicans*, suggesting that caspase-14–mediated IL-1β maturation in epithelial cells may represent a broad anti-bacterial immunity. Notably, caspase-14–mediated IL-1β maturation in epithelial cells dominates anti-*Y. pseudotuberculosis* IgG production, linking epithelial cell inflammation to anti-bacterial humoral immunity (**Extended Data Fig. 6**). Future studies will uncover other inflammasome/pyroptosis-targeting effectors and molecular/cellular mechanisms underlying the regulation of anti-bacterial humoral immunity by epithelial cell-derived IL-1β maturation.

It has been shown that *Y. pseudotuberculosis* effector protein YopJ blocks activation of transforming growth factor β–activated kinase 1 (TAK1), in host cells, thereby silencing cytokine expression ^21–23^. In response, host cells set off an alternative pyroptotic pathway, mediated by toll-like receptors (TLRs) or death receptors, which leads to the assembly of a complex that includes the FADD and the receptor-interacting serine-threonine protein kinase 1 (RIPK1) ^24,25^, which then recruits and activates caspase-8, which, in turn, cleaves the GSDMD protein to induce pyroptosis ^14–17^. Blockade of TAK1 with the small-molecule inhibitor, 5z-7-oxozeaenol (5z7), mimics the effect of YopJ and *Y. pseudotuberculosis* infection to cause pyroptosis in the presence of the TLR ligand, LPS, or inflammatory factor, tumor necrosis factor– α (TNF-α) ^14–16,22^. Activation of TAK1 and IKK blocks the function of caspase-8 by inducing the expression of cFLIP ^17^. Therefore, the function of caspase-14 on the maturation of IL-1β downstream of caspase-8 is likely to be inhibited during inflammasome activation, in which TAK1 and IKK is activated. This may explain why caspase-1, not caspase-14–mediated cleavage of IL-1β, is predominantly functional during inflammasome activation. However, blockade of TAK1 activation by a bacterial effector protein, such as YopJ, restores the function of caspase-8 as well as its downstream caspase-14–mediated processing of IL-1β. Interestingly, we and others observed the cleavage of caspase-1 to active caspase-1 p20 in response to *Y. pseudotuberculosis* or TNF plus 5z7 in WT macrophages ^14,16^. Even in *caspase-14*^-/-^ epithelial cells, active caspase-1 p20 was induced in response to *Y. pseudotuberculosis* or LPS plus 5z7. However, no detectable pro–IL-1β processing was observed in *caspase-14*^-/-^ epithelial cells, indicating that activation of TAK1/IKK may dictate the activity of caspase-1 active fragment toward the cleavage of pro-IL-1β. Through mass spectrometry analysis and *in vitro* verification, we identified that caspase-1 p20 Ser126 is directly phosphorylated by activated TAK1. Phosphorylation of caspase-1 at Ser126 by TAK1 is essential for its activation and subsequent cleavage of pro-IL-1β. Disruption of this phosphorylation of caspase-1 by TAK1 does not affect the processing of caspase-1, but abrogates its ability to cleave pro-IL-1β, suggesting that activation of caspase-1 may require two signals: one is for its cleavage into an active fragment, and another one is the phosphorylation of the active caspase-1 fragment by TAK1 to achieve its cleavage ability towards pro-IL-1β. However, pathogens like *Y. pseudotuberculosis* can utilize their effector protein such as YopJ to inhibit TAK1, thereby blocking caspase-1-mediated pro-IL-1β cleavage to evade host defense mechanisms. Unexpectedly, pathogen blockade of TAK1 restores the function of caspase-8 by inhibiting the expression of cFLIP ^17^, thus enabling host cells to set off an alternative caspase-14-dependent pathway to fulfill the cleavage and maturation of pro-IL-1β.

Classical DED-DED binding between caspase-8 and FADD triggers caspase-8 oligomerization and activation ^61,62^. Though caspase-14 does not possess a DED domain as caspase-8 does ^8,41^, it has a similar affinity with FADD as caspase-8 protein did *in vitro*. This binding promotes the dimmer formation and activation of caspase-14 p16. Using protein-protein docking analysis and experimental validation, we identified that caspase-14 residues E106 and Y162 interact with FADD DED domain residues K24 and S41. Unexpectedly, in the presence of caspase-8, while caspase-14’s FADD-interacting residues (E106/Y162) remained unchanged, the caspase-14-interacting sites on FADD switched from K24/S41 to S13/R72. These FADD sites (S13/R72) were essential for caspase-14 cleavage in LPS plus 5z7-stimulated A549 cells. These results suggest that distinct sites on FADD differentially mediate interactions with caspase-14 and caspase-8 within the complex, enabling proper caspase-14 cleavage and activation.

Previous studies established that a high concentration of the active caspase-8 fragment (300 ng) can directly cleave pro–IL-1β *in vitro* ^65^. Through *in vitro* examination, we found caspase-14 cleaves pro-IL-1β efficiently at low concentrations (30 ng), comparable to typical caspase-1 concentrations ^67^, suggesting caspase-14 as a predominant pro-IL-1β processor. Furthermore, macrophages from *Ripk3^-/-^Casp8^-/-^* or no-cleavable *Casp8^D387A/D387A^* mice exhibit deficient IL-1β cleavage ^26–29^. In our study, we found that caspase-8 deficiency in LPS/5z7-stimulated A549 cells or epithelial cells abolished both caspase-14 and IL-1β cleavage. In contrast, caspase-14 deficiency specifically blocked pro-IL-1β cleavage while preserving caspase-8 activation in both A549 and epithelial cells. Furthermore, *Y. pseudotuberculosis* infection or LPS plus 5z7 stimulation failed to induce the cleavage of pro–IL-1β in *caspase-14*^-/-^ A549 or epithelial cells complemented with a caspase-14 (D146A) mutant, which is resistant to the cleavage by caspase-8. Collectively, these findings indicate that caspase-8 mediates the cleavage and activation of caspase-14, which subsequently processes pro-IL-1β under physiological conditions in epithelial cells.

Caspase-11 serves as the essential sensor and mediator for cytoplasmic LPS-induced non-canonical pyroptosis ^12,13,38,82^. Its interdependent activation with NLRP3 drives the cleavage of caspase-1, GSDMD, and IL-1β, culminating in pyroptosis and cytokine release ^83,84^. However, LPS electroporation induced no detectable caspase-14 cleavage in either WT or *Casp-1*^−/−^ *Casp-11*^−/−^primary epithelial cells. Furthermore, caspase-1 or caspase-11 deficiency did not impair caspase-14 or pro-IL-1β cleavage in LPS plus 5z7-stimulated primary epithelial cells. These results suggest caspase-14 functions independently of the caspase-11 pathway in regulating IL-1β maturation.

Though the molecular and cellular function of caspase-14 is poorly characterized, the majority of clinical data indicate the role of caspase-14 in the etiopathology of many diseases such as retinal dysfunctions, multiple malignancies, and skin conditions ^45,85,86^. Deletions in the human *CASP14* gene have been linked to a defect in cornification that manifests as autosomal recessive inherited ichthyosis ^87^. Unlike other caspases, caspase-14 has a restricted expression pattern in the embryo and skin ^8,88^. Furthermore, caspase-14 does not participate in apoptotic pathways but its processing is associated with epidermal differentiation ^45,85^. We found that caspase-14 acts as a novel regulator of IL-1β maturation downstream of caspase-8–mediated pyroptosis. Given that IL-1β plays a key role in a variety of diseases, whether and how caspase-14–mediated processing and maturation of IL-1β is involved in other types of related diseases in response to different stresses or stimuli needs further investigation.

## Methods

### Bacterial strains and cells

The *Legionella pneumophila*, a gift from Dr. Yongqun Zhu (Center for Veterinary Sciences, Zhejiang University, Hangzhou, China); *Y. pseudotuberculosis* YPIII strain, a gift from Dr. Shiyun Chen (Wuhan Institute of Virology, Chinese Academy of Sciences), was grown overnight in 2× YT broth at 26°C. On the day of infection, bacteria were diluted 1:50 into 2× YT plus 20 mM MgCl_2_ and 20 mM sodium oxalate and grown for 2 hours at 26°C, followed by a shift to 37°C for 2 hours. Bacteria were then washed in phosphate-buffered saline (PBS; Invitrogen) and added to cells at a MOI of 40. Next, 100 μg/ml gentamicin was added to the cultures 2 hours after infection. To quantify the number of bacteria that had been taken up by cells, iBMDMs were infected with the *Y. pseudotuberculosis* YPIII strain at the MOI indicated in the corresponding figures. Thirty minutes later, cells were washed with PBS three times and gentamicin was added to kill extracellular bacteria. Then, intracellular bacteria were released by treating cells with 0.05% Triton X-100 before lysates were serially diluted and plated on 2× YT agar. Bacterial colonies were counted after 1 day of culture at 37°C.

A549 cells (ATCC TCH-C116) and HEK293T cells (ATCC CRL-3216) were resuspended in DMEM (HyClone) mixed with 10% (v/v) fetal bovine serum (FBS, Gibco) for experiments. Macrophages were cultured in RPMI-1640 medium supplemented with 10% (v/v) FBS. Peritoneal macrophages were obtained from mice(4∼6weeks) three days after injection of thioglycollate (BD). All the cells were routinely tested for contamination by mycoplasma.

### Y. pseudotuberculosis infection

On the day of infection, bacteria were diluted 1:50 into LB broth and grown for 4 hours at 37°C. Bacteria were then washed with PBS and added to cells at a MOI of 20. Next, 100 μg/ml gentamicin was added to the cultures 0.5 hours after infection to kill extracellular bacteria.

For in vivo infection, mice were fasted for 16 hours and challenged with 1 × 10^7^ bacterial burdens, mice were euthanized related days after infection and tissues were harvested, homogenized in 1 ml of phosphate-buffered saline (PBS), and serially diluted on LB agar.

### Caspase-14-Knockout Mice

C57BL/6J Cya-Casp14^em1flox/Cya^ (S-CKO-18523) C57BL/6J-*Col1a2*^em1(P2A-iCre),^ (*Col1a2-iCre*) (Cat. C001528),C57BL/6JCya-Ccsp^em1(P2A-iCre)/Cya^(Cat.I001145), C57BL/6JCya-Igs2em1^(Vil1-MerCreMer)^/Cya (Cat. C001433), C57BL/6JCya-Casp14^em1/Cya^ mice were purchased from Cyagen. To obtain conditional Caspase-14-KO mice fibroblast (for short as *Casp-14^f/f^ Col1a2^Cre^*), epiblasts cells of lung (for short as *Casp-14^f/f^ Ccsp^Cre^*) or epiblasts cells of intestine (for short as *Casp-14^f/f^Villin^Cre^*), 4-week-old *Caspase-14*^flox/flox^ female mice were mated with male C57BL/6J-*Col1a2*^em1(P2A-iCre)^,(*Col1a2-iCre*) (Cat. C001528) or C57BL/6JCya-Ccsp^em1(P2A-iCre)/Cya^ (Cat.I001145), C57BL/6JCya-Igs2em1^(Vil1-MerCreMer)^/Cya (Cat. C001433) mice. C57BL/6JCya-Lyz2^em2(IRES-iCre)/Cya^ (Cat.C001358) was used for *^-^* whole-body gene knockout mice. Moreover, all animal experiments were reviewed and approved by the Animal Experiment Administration Committee of XX(Tongji) University School of Medicine.

### Caspase-14^C136A/C136A^ or Caspase-1^D156A/D156A^ knockin mice

The procedure for the generation of *Caspase-14^C136A/C136A^*or *Caspase-1^D156A/D156A^* knockin mice is similar to our previous report. The T7 promoter sequence was fused with the Cas9 coding region which was cloned from pX260 plasmid (Addgene #42229). Similarly, T7 promoter sequence and the targeting sequence of Caspase-14 were fused to the guide RNA scaffold which was cloned from pX330 plasmid (Addgene # 42230). In vitro transcription of Cas9 mRNA and sgRNAs targeting STING was performed with mMESSAGE mMACHINE T7 ULTRA kit

(ThermoFisher Scientific, AM1345, Waltham, MA) and MEGAshortscript T7 kit (ThermoFisher Scientific, AM1354, Waltham, MA) according to the manufacturer’s instructions, respectively. Both Cas9 mRNA and sgRNAs were then purified using MEGAclear Transcription Clean-Up Kit (ThermoFisher Scientific, AM1908, Waltham, MA) and stored at −80°C.

One-cell embryos were collected from superovulated wild type C57BL/6J female mice that had been mated with wild type C57BL/6J male mice. Cas9 mRNA (50 ng/μL), sgRNA targeting Caspase-14 (50 ng/μL) and donor (ssODN, 100 ng/μL) were mixed together and then injected into the one-cell embryos. The injected embryos were cultured in EmbryoMax KSOM Medium (Sigma-Aldrich, MR-106-D, USA) until the two-cell stage, followed by transferring into oviducts of recipients at 0.5 dpc. Recipient female mice delivered pups at 19.5 dpc. The first generation of point-mutant mouse were identified using the genotyping primers. The mice harboring the correct point mutation were crossed for expansion of the mouse population.

### Isolation of cells from mouse lungs or intestine

The isolation of pulmonary epithelial cells was performed via enzymatic digestion and physical disaggregation. Briefly, lung tissues were subjected to digestion in DMEM supplemented with 200 U ml−1 Collagenase I (Gibco), 5 Unit ml−1 Dispase (Worthington), 0.1 mg ml−1 DNase I (Sigma-Aldrich), and 4 Unit ml−1 Elastase (Worthington) for 45 minutes at room temperature. The digested tissue homogenate was then filtered through a series of 70-μm and 40-μm cell strainers to obtain a single-cell suspension. After erythrocyte lysis, the target population of CD45-negative and EpCAM-positive (CD45−EpCAM+) lung epithelial cells was isolated using fluorescence-activated cell sorting (FACS).

The isolation of colonic epithelial cells was performed using a standard protocol based on chelation and mechanical dissociation. Briefly, following dissection, the colons were thoroughly cleansed with phosphate-buffered saline (PBS) and sectioned into small segments. These segments were then subjected to incubation in Hank’s Balanced Salt Solution (HBSS) containing 5 mM ethylenediaminetetraacetic acid (EDTA) and 0.5 mM dithiothreitol (DTT) at 37°C for 30 minutes with constant gentle agitation. This process helps dissociate the epithelial layer from the underlying tissue. The resulting supernatant, which contains the sloughed-off epithelial cells, was subsequently passed through a 70 μm cell strainer to remove large aggregates and debris, followed by two washing steps. The purified epithelial cells were then lysed using a buffer containing 1% sodium dodecyl sulfate (SDS). The lysates were clarified via sonication until a clear solution was obtained, making them suitable for subsequent western blot analysis

### Caspase-8 KO, GSDMD KO, and FADD KO cells

LentiCRISPRv2 vectors were utilized to generate knockout (KO) cells. HEK293T cells were transfected by mixture of Lipofectamine 2000 (Invitrogen, 11668030) with pMD2.G, pSPAX2 and LentiCRISPRv2 (Changsha Youbio Tech, VT8107) harboring the guide (g)RNA that targeted TTCCACTTCTACGA-TGCCA for GSDMD, GCTCTTCCGAATTAATAGAC for Caspase-8, TTCCTATGCCTCGGGCGCGT for FADD, GGTGTTGATGAGCTGCGCGG for Caspase-1,

GGTGTTGATGA-GCTGCGCGG for Caspase-11 in A549 cells or scrambled gRNA. The lentiviruses were harvested 48 h post transfection and were utilized to infected cells. The infected cells were then selected with puromycin (4 μg/mL) and single cell clone was acquired through serial dilutions in a 96-well plate. The KO cells were confirmed by Western blot and used for further experiments.

### Reagents, and antibodies

The following antibodies and reagents were used for western blot, IF or CO-IP: CASP14 Polyclonal antibody (28136-1-AP), CD107b / LAMP2 Monoclonal antibody(66301-1-Ig), CoraLite® Plus 488 Anti-Mouse CD107a/LAMP1 (1D4B) (CL488-65050), CoraLite® Plus 488-conjugated Caspase-8/p43/p18 Monoclonal antibody (CL488-66093), CoraLite® Plus 647-conjugated Phospho-RIPK1 (Ser161) Monoclonal antibody (CL647-66854) Caspase 1/P20 Polyclonal antibody (22915-1-AP) were from Proteintech. Monoclonal anti-Flag antibody (F3165), and anti–β-actin antibody (A1978) were from Sigma-Aldrich. Antibodies against HA (#3724, #2367), Flag (#14793), cleaved Caspase-8 (#8592), Caspase-8 (#4927), RIPK1 (#3493), Phospho-RIPK1 (#31122) were from Cell Signaling Technology. Monoclonal anti-FADD antibody (sc-166516) was from Santa Cruz Biotechnology. Monoclonal anti-GSDMD antibody (ab209845), monoclonal anti–pro Caspase-8 antibody (ab108333) and monoclonal anti-FADD antibody (ab124812) were from Abcam.LPS (#L4524), etoposide (#E1383), 5z-7 (O9890) were from Sigma-Aldrich. Recombinant murine TNF-α (#315-01A) was from Peprotech. Nigericin (tlrl-nig) were from Invitrogen.

### Generation of phospho-Caspase-1^S1^^26^ antibody

Rabbit polyclonal antibody was raised against Caspase-1 phosphorylated on S126 (p-Caspase-1^S1^^26^) in collaboration with Abclonal Biotech. In brief, 3 rabbits were immunized with peptide c(KLH)-PT(S-p)-SGSEGN (in which ‘c(KLH)’ indicates keyhole limpet haemocyanin fused through cysteine, and ‘p-S’ indicates phosphorylated serine) at a ratio of 1:1. The nonphosphorylated peptide c(KLH)-PT-S-SGSEGN was used for antibody purification and detection.

### ATP loss assay

For the CellTiter-Glo assay, CTG reagent (Promega, G7570) was mixed at a 1:1 ratio with supernatant from the treatment plate. The mix was incubated for 10 min at room temperature on the shaker, followed by luminescence measurement.

On the day of infection, bacteria were diluted 1:50 into LB broth and grown for 4 hours at 37°C. Bacteria were then washed with PBS and added to cells at a MOI of 20. Next, 100 μg/ml gentamicin was added to the cultures 0.5 hours after infection to kill extracellular bacteria.

### Genome-wide CRISPR-Cas9 screen

The Mouse_GeCKOv2_Library_A/B was obtained from AZENTA and amplified according to the protocol provided by the manufacturer. For lentivirus production, mouse GeCKO A and B library plasmids were mixed equally and transfected into HEK293T cells together with the packaging plasmids psPAX2 and pMD2.G, Seventy-two hours later, lentivirus was collected and the viral titer was measured with the QuickTiter Lentivirus Titer kit (Cell Biolabs, #VPK-107). For the genome-wide screen, Cas9 stably expressing WT or *Caspase-1^-/-^11^-/-^* A549s were seeded in 10-cm dishes (2 × 106 cells/dish), and 7 × 107 cells were infected with the lentivirus-containing sgRNA library at a multiplicity of infection (MOI) of 0.3. Sixty hours later, cells were treated with puromycin to remove uninfected cells. Six days after that, the transduced cells were seeded in 40 × 10 cm dishes (8 × 106 cells/dish). Plasmids eGFP-IL-1β-mCherry was overexpressed in cells and Cells both GFP and mCherry expression were sorted expanded in DMEM supplemented with 10% FBS. Expanded cells were treated with 10 ng/ml TNF-α plus 125 nM 5z7 for 3 hours and cells lacking mCherry expression were sorted and expanded in DMEM supplemented with 10% FBS. Expanded cells and untreated transduced cells (as the control sample) were harvested and lysed in the SNET buffer [20 mM Tris-HCl pH 8.0, 5 mM EDTA, 400 mM NaCl, 1% SDS and 400 μg/ml Proteinase K (NEB, P8107S)]. Genomic DNAs were prepared by using phenol-chloroform extraction and isopropanol precipitation and amplified by two-step PCR using the 2× Hieff Canace Gold PCR Master Mix (Yeasen, #10149ES01). The samples were quantified and sequenced on a HiSeq 2500 (Illumina) by GENEWIZ. Sequencing data were further processed and analyzed using the MAGeCK algorithm. MAGeCK built a linear model to estimate the variance of guide RNA (gRNA) read counts, evaluated the gRNA abundance changes between control and treatment conditions, and assigned P values for positive and negative selection.

### Lysosome extraction

In all, 1 × 10^8^cells per sample were grown to 90% confluency, treated with TNF-α and 5z-7. All subsequent steps of the lysosomal isolation were performed according to manufacturer’s description (LYSISO1, 233-140-8). In brief, cells were centrifuged at 600 ×*g*for 5 min, resuspended in 2.7 packed cell volume of 1 × extraction buffer and homogenized in a glass Dounce homogenizer. The nuclei were removed by centrifugation at 1.000 ×*g* for 10 min. The postnuclear supernatant was centrifuged at 20.000 ×*g*for 20 min and the resulting pellet, containing the crude lysosomal fraction, was resuspended in a minimal volume of 1 × extraction buffer (0.4 ml per 10^8^cells). To enrich the lysosomes, the suspension was further purified by density gradient centrifugation at 150.000 ×*g*for 4 h on a multistep OptiPrep (Sigma-Aldrich, Steinheim, Germany) gradient according to the manufacturer’s description. Altogether, 0.5 ml fractions were collected starting from the top of the gradient. Each fraction was assayed for Caspase-8, GSDMD and Lamp2 by western blotting.

### Mice immunisation and ELISA

Female C57BL/6 (6 weeks old) were immunized via aerosol infected with bacterial strains resuspended in 200 μl of PBS. The immunization doses were adjusted based on the virulence of each strain, specifically using 1 × 10^6^ CFU for the wild-type *Y. pseudotuberculosis.* Serum samples were collected 10 days post-immunization, and *Y. pseudotuberculosis* -specific antibody levels were quantified by a standard indirect ELISA. Briefly, 96-well plates were coated overnight at 4°C with 2×10^7^ CFU of live *Y. pseudotuberculosis* carbonate-bicarbonate buffer. After washing with PBST, plates were blocked with 2% BSA in PBST for 2 hours at 37°C. Serial twofold dilutions of mouse sera were added to the plates and incubated for 2 hours at 37°C, followed by incubation with an HRP-conjugated goat anti-mouse IgG antibody (1:6,000 dilution) for 1 hour at 37°C. The reaction was developed using o-phenylenediamine substrate for 15 minutes at room temperature and stopped with 2 M H SO . The optical density was measured at 450 nm.

### Analysis of antibody effects on *Y. pseudotuberculosis* infection in vivo

C57BL/6J mice were tail vein injected by normal mouse IgG or IgG isolated from *Y. pseudotuberculosis* infected WT or conditional knock-out Caspase-14 in lung epithelial cells or knock-in Caspase-14C136A mice (*Y. pseudotuberculosis* WT/Il-1b^-/-^/Casp-14 KO/ KI IgG) (15 mg/kg mouse) for 24 hours followed by aerosol injected with 1× 10^7^ CFU of *Y. pseudotuberculosis*. Normal mouse IgG was purchased from Beyotime (A7028). 3 days after *Y. pseudotuberculosis* infection, the mice were sacrificed, the lung removed, and the number of *Y. pseudotuberculosis* CFUs were monitored.

### Western blot

For Immunoblot, cell lysates or precipitates in 1× sodium dodecyl sulfate (SDS) protein sample buffer were denatured at 95 °C for 8 min and then were resolved by electrophoresis through a 4% to 15% SDS-polyacrylamide gel. Separated proteins were transferred onto polyvinylidene difluoride (PVDF) membranes and were blocked with 5% skimmed milk for more than one hour. The membranes were then incubated with the prespecified antibodies at the indicated dilutions. An enhanced chemiluminescence reagent (Thermo Fisher Scientific) was applied for Immunoblot.

### Immunoprecipitation assay

Cells were lysed using RIPA Lysis Buffer (Beyotime Biotechnology, China) supplemented with protease inhibitor cocktail (P8340, Sigma-Aldrich), 1 mM of PMSF and phosphatase inhibitor cocktail (P5726, Sigma-Aldrich). The lysates were centrifuged at 12,000 rpm for 10 min and the cellular debris was discarded. For immunoprecipitation, cell lysates were incubated with monoclonal anti-HA agarose (A2095, Sigma-Aldrich), Anti-FLAG M2 Affinity Gel (A2220, Sigma-Aldrich), or corresponding antibodies at 4°C overnight. The immunoprecipitations were subjected for further analysis.

### Analysis of Protein docking Prediction Results

In this study, the professional HDCOK program for protein-protein and protein-DNA/RNA docking was adopted for docking. This docking was a global docking. After the docking was completed, the structure with the best docking score was selected as the standard result for subsequent interaction analysis. The connection score is calculated based on the ITScorePP or ITScorePR iterative scoring function. When the docking score is negative, the larger its absolute value is, the greater the possibility of the combination of this combined model and the stronger the interaction. Take PDB into consideration, the docking score of the protein-protein complex in it is usually around -200 or lower. Based on experience, we have defined a confidence score dependent on the docking score to indicate the possibility of binding between the two molecules As shown below: Confidence_score = 1.0/[1.0+e0.02*(Docking_Score+150)]

### Immunofluorescent assay, confocal microscopy

Cells were fixed with 4% paraformaldehyde (PFA) in PBS for 25 min at room temperature. Cells then were blocked and permeabilized for 30 min in blocking buffer (PBS containing 1% bovine serum albumin (BSA) and 0.1% Triton X-100). Cells were then incubated with primary antibody in 4 °C for overnight and secondary antibody for 1 h at room temperature. Nuclei were labelled by staining with DAPI. Images were collected using a Leica TCS SP8 confocal laser microscopy system (Leica Microsystems, Buffalo Grove, IL) and processed with ImageJ (V1.8.0.112). Manders’ overlap coefficient was calculated using ImageJ (where each point represents 30 cells). All images are representative of at least three independent experiments.

### Mass spectrometry analysis

Analysis of caspase-1 phosphorylation sites Flag-caspase-1 stably expressed HEK293T cells that transfected with HA-TAK1 or HA-TAK1 together with HA-TAB1 was immunoprecipitated with anti-caspase-1 antibody. The beads were then washed and boiled into 1× SDS loading buffer for SDS-PAGE analysis, followed by Commassie blue staining. The caspase-1 protein bands were destained, reduced and alkylated and then digested with trypsin. The peptides were extracted with 0.5% formic acid and 50% acetonitrile followed by 0.1% formic acid and 80% acetonitrile. The extracted peptides were bound to a Magic C18 AQ reverse phase column (100 μm × 50 mm; Michrom Bioresources). An Agilent 1100 binary pump was used to generate the HPLC gradient as follows: 0–5% B for 5 min, 5–45% B for 40 min, 45–80% B for 3 min, and then to 5%B in 2 min (A = 0.1 M acetic acid in water; B = 0.1 M acetic acid and 70% acetonitrile). The eluted peptides were sprayed into an LTQ Orbitrap XL mass spectrometer (Thermo Electron Corp) that was equipped with a nano-ESI ion source. The mass spectrometer was operated in data-dependent mode with automatically switch between mass spectrometry and tandem mass spectrometry acquisition. The resulting MS/MS data were processed using Proteome Discoverer 1.4 (ThermoFisher). Tandem mass spectra were searched against human caspase-1 protein sequence. Trypsin/P was specified as cleavage enzyme allowing up to 2 missing cleavages. Mass error was set to 20 ppm for precursor ions and 0.05 Da for fragments ions. Carbamidomethyl on Cys were specified as fixed modification, oxidation on Met and phosphorylation on Ser and Thr were specified as variable modification. Peptide confidence was set at high, and peptide ion score was set >20. The mass spectrometry analysis was performed by JingJie PTM Biolabs (Hangzhou, China).

### *In vitro* kinase assay

The *In vitro* kinase assay was performed as reported previously. 2 μg recombinant caspase-1 or caspase-1 mutant protein was incubated with 2 μg recombinant TAK1 proteins or TAK1/TAB1 complex in 1× kinase assay buffer (CST, #9802) for 30 min at 30°C. Then the reaction system was diluted with RIPA lysis buffer. The protein samples were then resolved by electrophoresis through a 10% SDS-polyacrylamide gel and further applied for immunoblot.

### Recombinant GST-IL-1β cleavage in vitro assay

Recombinant GST-IL-1β (5ug) was incubated in the presence or absence of 5ug recombinant Caspase-8, recombinant protein Caspase-14 or FADD Fusion Protein (Proteintech) at 37 °C for 60 min in Caspase reaction buffer (20 mM PIPES pH 7.2, 10% sucrose, 150 μM ATP,100 mM NaCl, 0.1% CHAPS, 1 mM EDTA, 10 mM DTT) with or without Kosmotropic salts (1.1 M sodium citrate). Samples were then subjected to SDS-PAGE.

### Statistical analysis

Statistical significance between groups was determined by two-tailed Student’s t-test, two-tailed analysis of variance followed by Bonferroni post hoc test or two-sided Mann–Whitney U-test. Differences were significant at P < 0.05. The experiments were not randomized, and the investigators were not blinded to allocation during experiments and outcome assessment.

## Author contributions

Conceptualization:
Methodology:
Investigation:
Funding acquisition:
Supervision:
Writing:

## Data and materials availability

The main data supporting the findings of this study are available within the paper. Additional data are available from the corresponding authors upon reasonable request. This study did not generate new unique reagents but processed data generated for this study can be found in the supplementary files.

