## Extended data for "Caspase-14 recognizes and processes IL-1β in epithelial cells to drive anti-bacterial IgG production"

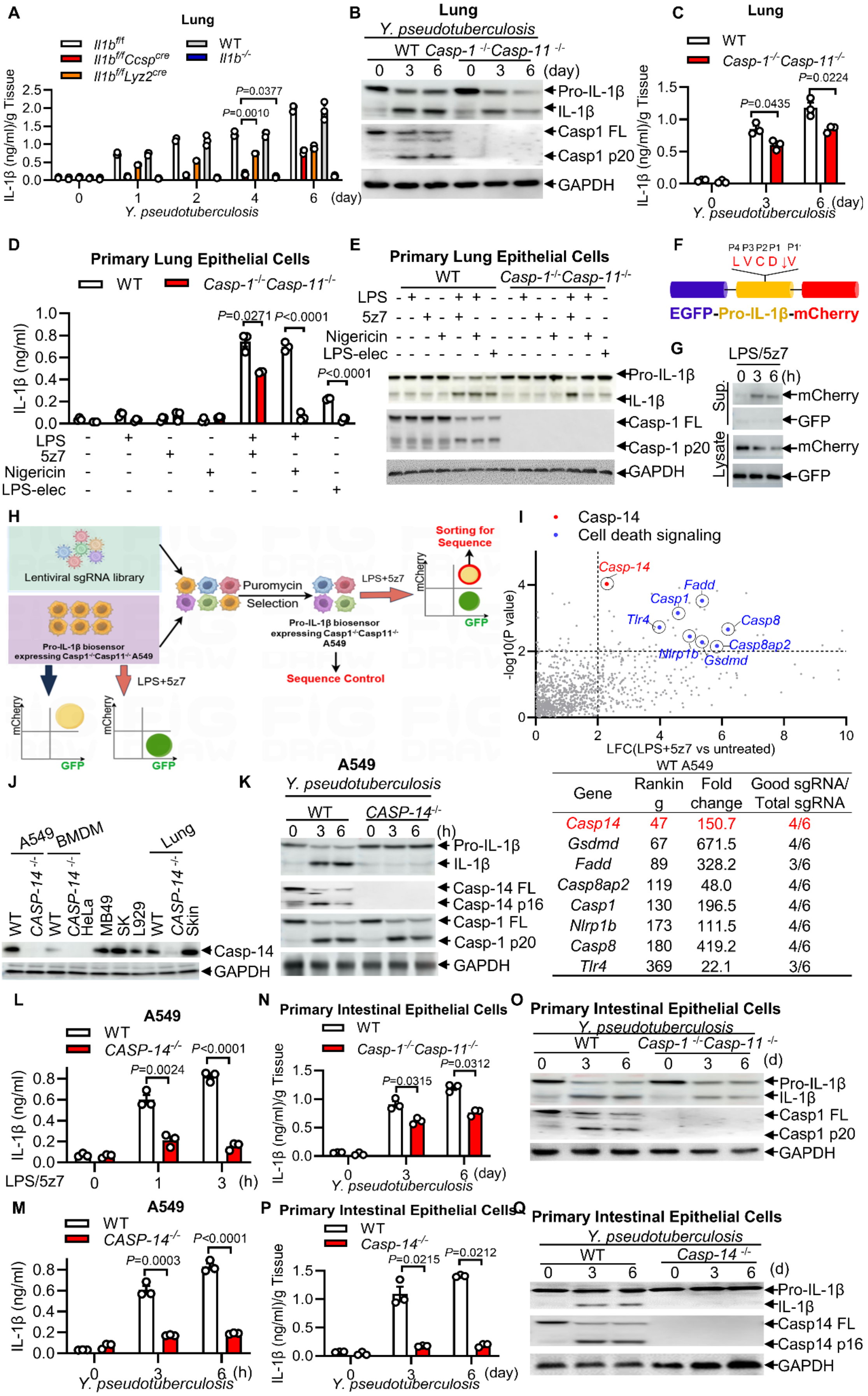


WT A549

| Gene | Ranking | Fold change | Good sgRNA/ Total sgRNA |
|---|---|---|---|
| *Casp14* | 47 | 150.7 | 4/6 |
| *Gsdmd* | 67 | 671.5 | 4/6 |
| *Fadd* | 89 | 328.2 | 3/6 |
| *Casp8ap2* | 119 | 48.0 | 4/6 |
| *Casp1* | 130 | 196.5 | 4/6 |
| *Nlrp1b* | 173 | 111.5 | 4/6 |
| *Casp8* | 180 | 419.2 | 4/6 |
| *Tlr4* | 369 | 22.1 | 3/6 |

**Extended Data Fig. 1. Caspase-14 regulates maturation of IL-1β**

**(A)** 6-8 weeks female *Casp-14*$^{f/f}$ or *Casp-14*$^{f/f}$ *Ccsp*$^{Cre}$ for epithelial cells, *il-1β*$^{f/f}$ *Lyz2*$^{Cre}$ for macrophages or *il-1b*$^{-/-}$ whole-body gene knockout mice orally challenged with *Y. pseudotuberculosis* (~$10^7$ CFU/mice). IL-1β release serum for indicated days post infection. **(B)** Immunoblotting of lung from WT or *Caspase1*$^{-/-}$ *Caspase-11*$^{-/}$ mice stimulated with *Y. pseudotuberculosis* for the indicated times. **(C)** IL-1β release was measured by ELISA with *Y. pseudotuberculosis* challenged WT*, Caspase1*$^{-/-}$*Caspase-11*$^{-/-}$ mice for the indicated times. **(D)** IL-1β release of epithelial cells was measured by ELISA with LPS electroporation, LPS/nigericin, LPS with or without 5z-7 -challenged for the indicated times. **(E)** Immunoblotting of epithelial cells from WT or *Caspase1*$^{-/-}$ *Caspase-11*$^{-/}$ mice stimulated with LPS electroporation, LPS/nigericin, LPS with or without 5z-7 for the indicated times. **(F, G)** Detecting multiple time points of IL-1β cleavage by FACS Schematic workflow of the genome-wide CRISPR screening procedure in A549 **(H, I)** List of sgRNA hits from the screening relative to controls in positive selection. The dotted lines represent P = 0.01. List of sgRNA hits from the screening in WT A549 treated with LPS/5z-7. Corresponding genes targeted with multiple sgRNAs and fold enrichments are shown. FACS Schematic workflow identify epithelial cells from WT or *caspase-14*$^{-/-}$ mice **(J)** Immunoblotting of indicated proteins different cells. **(K)** Immunoblotting of indicated proteins from *WT* or *Caspase14*$^{-/-}$ of A549 stimulated with *Y. pseudotuberculosis,* LPS electroporation, LPS/nigericin or LPS with or without 5z-7 for the indicated times. **(L, M)** IL-1β release was measured by ELISA from *WT or Caspase14*$^{-/-}$ of epithelial cells from intestine stimulated with *Y. pseudotuberculosis* or LPS with or without 5z-7. (**N, P)** IL-1β release was measured by ELISA. (**O, Q)** Immunoblotting of indicated proteins from *WT or Caspase14*$^{-/-}$ of epithelial cells from intestine stimulated with *Y. pseudotuberculosis*. All of the immunoblot data are representative images from one of three independent experiments. Results in **A, C, D, L, M, N** and **P** reflect the mean ± s.e.m from three independent biological experiments. Two-tailed unpaired Student's t-test were used.

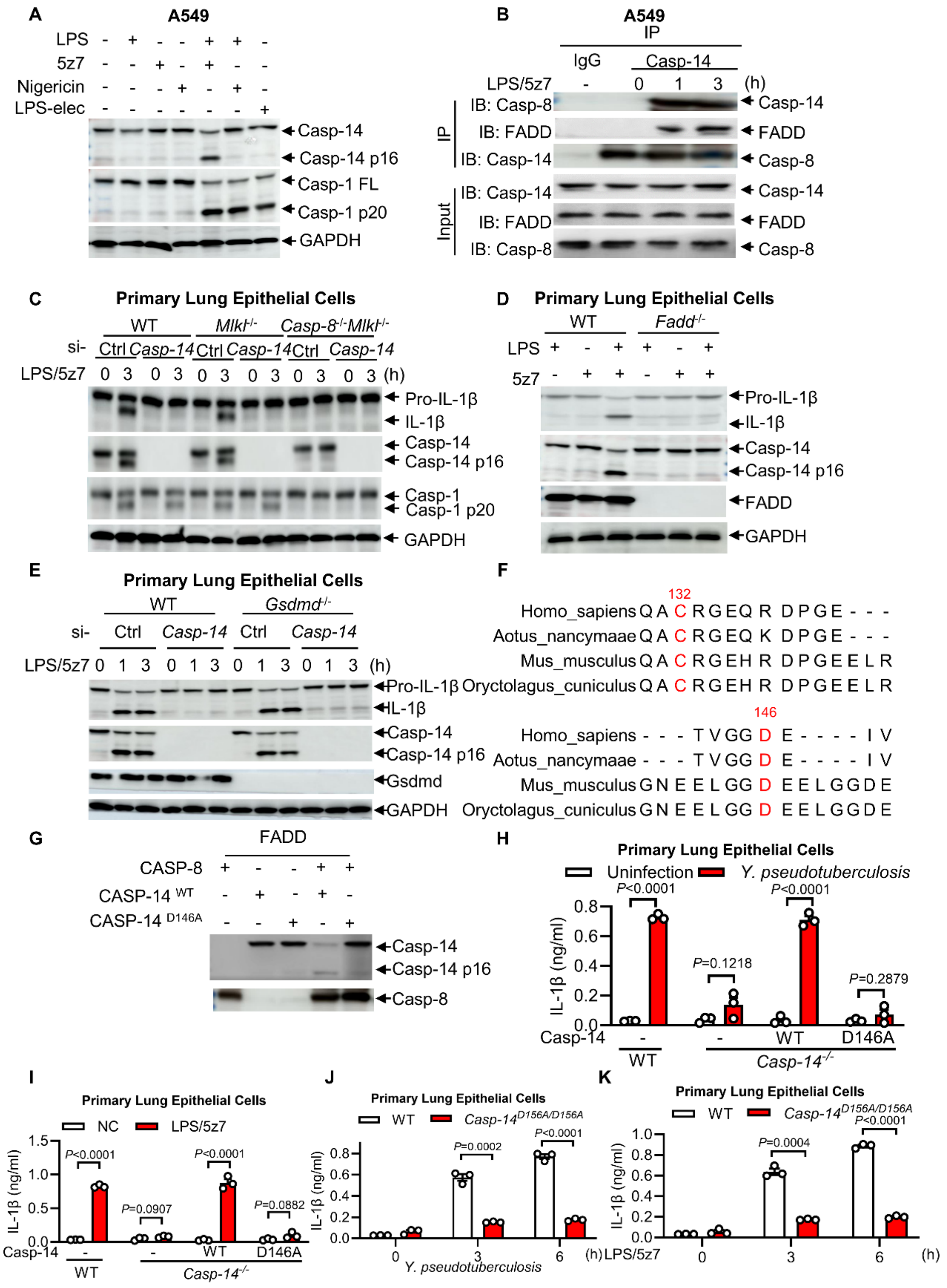


**Extended Data Fig. 2. Caspase-8 activates caspase-14**

**(A)** Immunoblotting of indicated proteins from A549 stimulated with LPS electroporation, LPS/nigericin or LPS with or without 5z-7. **(B)**Endogenous caspase-8 complex was immunoprecipitated with anti-caspase-8 antibody and analyzed by immunoblot with the indicated antibodies. **(C-E)** Immunoblotting of indicated proteins in from epithelial cells of WT *Caspase-8*$^{-/-}$ *MLKL*$^{-/-}$ , *FADD*$^{-/-}$, *GSDMD*$^{-/-}$ stimulated with LPS/5z-7 combined with si-*NC* or si-*Caspase-14* for the indicated times. **(F)**

Sequence alignment of conservation of the region of Caspase-14 harboring C132 and D146 amongst mouse, rat, human and monkey species. **(G)** In vitro cleavage assay of Caspase-14 or caspase-14 D146A protein with or without Caspase-8. **(H-K)** IL-1β release was measured by ELISA epithelial cells of WT *caspase14*$^{-/-}$ or *caspase-14* $^{D156A/D156A}$ *KI* stimulated with LPS/5z-7. All of the immunoblot data are representative images from one of three independent experiments. Results in **H-K** reflects the mean ± s.e.m from three independent biological experiments. Two-tailed unpaired Student's t-test were used.

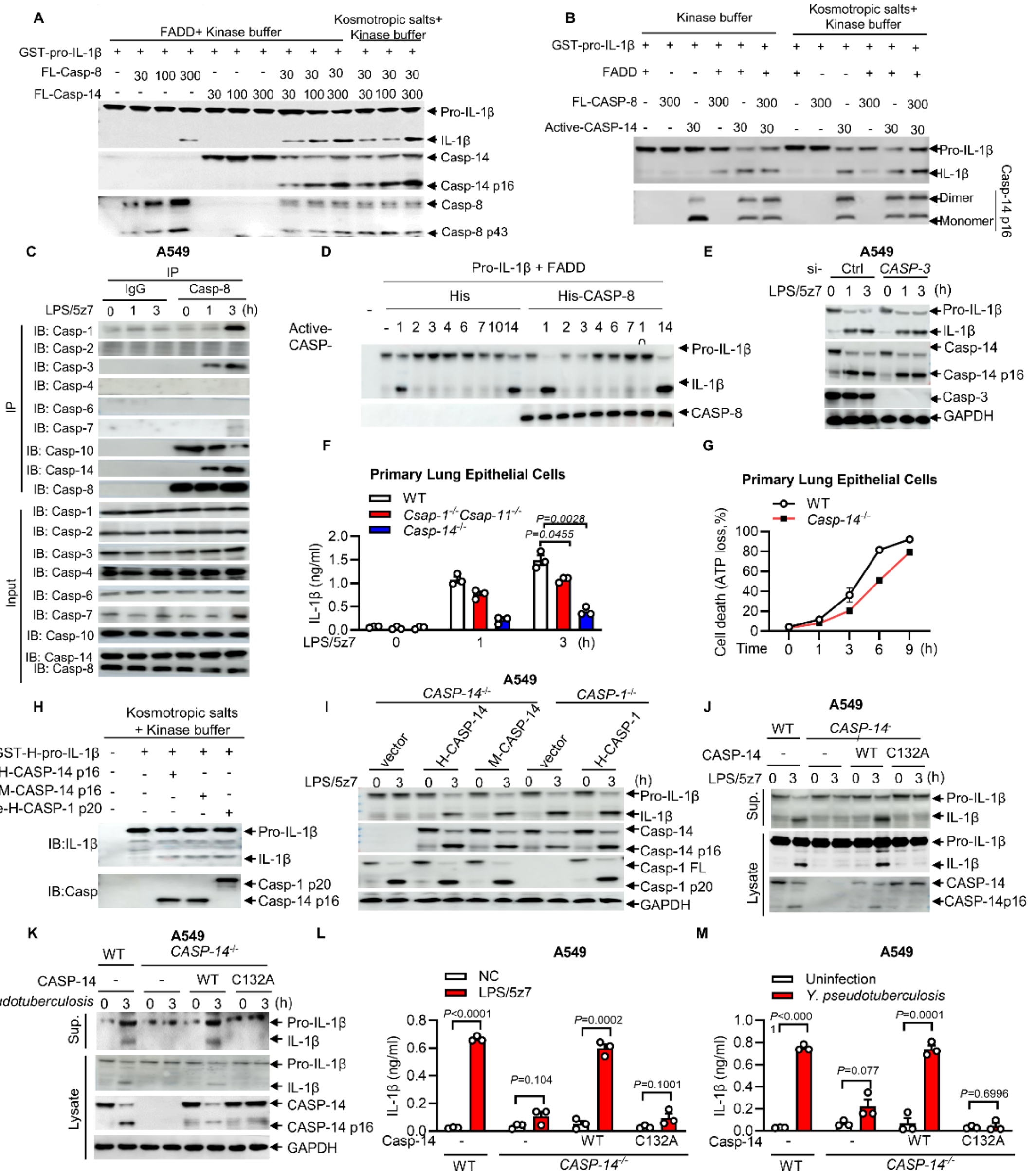


**Extended Data Fig. 3. Processing of pro–IL-1β by caspase-14 p16**

**(A, B)** In vitro cleavage assay of IL-1β protein with indicated proteins (30,100,300ng) combined with FADD or not in cleavage kinase buffer with or without Kosmotropic salts (1.1 M sodium citrate). **(C)** Endogenous caspase-8 complex was immunoprecipitated with anti-caspase-8 antibody and analyzed by immunoblot with the indicated antibodies. **(D)** In vitro cleavage assay of IL-1β protein with indicated active caspase proteins (30ng) combined with caspase-8(30ng) or not in cleavage kinase buffer. **(E)** Immunoblotting of indicated proteins from si-NC or si-Caspase3 A549 stimulated with LPS+ 5z-7 for the indicated times. **(F)** IL-1β release was measured by ELISA from WT, *Caspase1*$^{-/-}$or *Caspase14*$^{-/-}$ of A549 stimulated with LPS/5z-7. **(G)** WT and *Caspase-14*$^{-/-}$ A549 treated stimulated with LPS+ 5z-7. Cell death was assessed and calculated by measuring the ATP level at indicated times. **(H)** In vitro cleavage assay of IL-1β protein with human or murine caspase-14 or caspase-1 active in cleavage kinase buffer with Kosmotropic salts (1.1 M sodium citrate). **(I)** Immunoblotting of indicated proteins from *caspase14*$^{-/-}$ stably expressed human or caspase-14 WT or mutants stimulated with *Y. pseudotuberculosis* or LPS + 5z-7 for the indicated times. **(J, K**) Immunoblotting of indicated proteins from epithelial cells of WT or *Caspase14*$^{-/-}$ stimulated with *Y. pseudotuberculosis* or LPS/5z-7 for the indicated times. **(L, M)** IL-1β release was measured by ELISA stimulated with *Y. pseudotuberculosis,* or LPS/5z-7. All of the immunoblot data are representative images from one of three independent experiments. Results in **F, G, L, M** reflects the mean ± s.e.m from three independent biological experiments. Two-tailed unpaired Student's t-test were used.

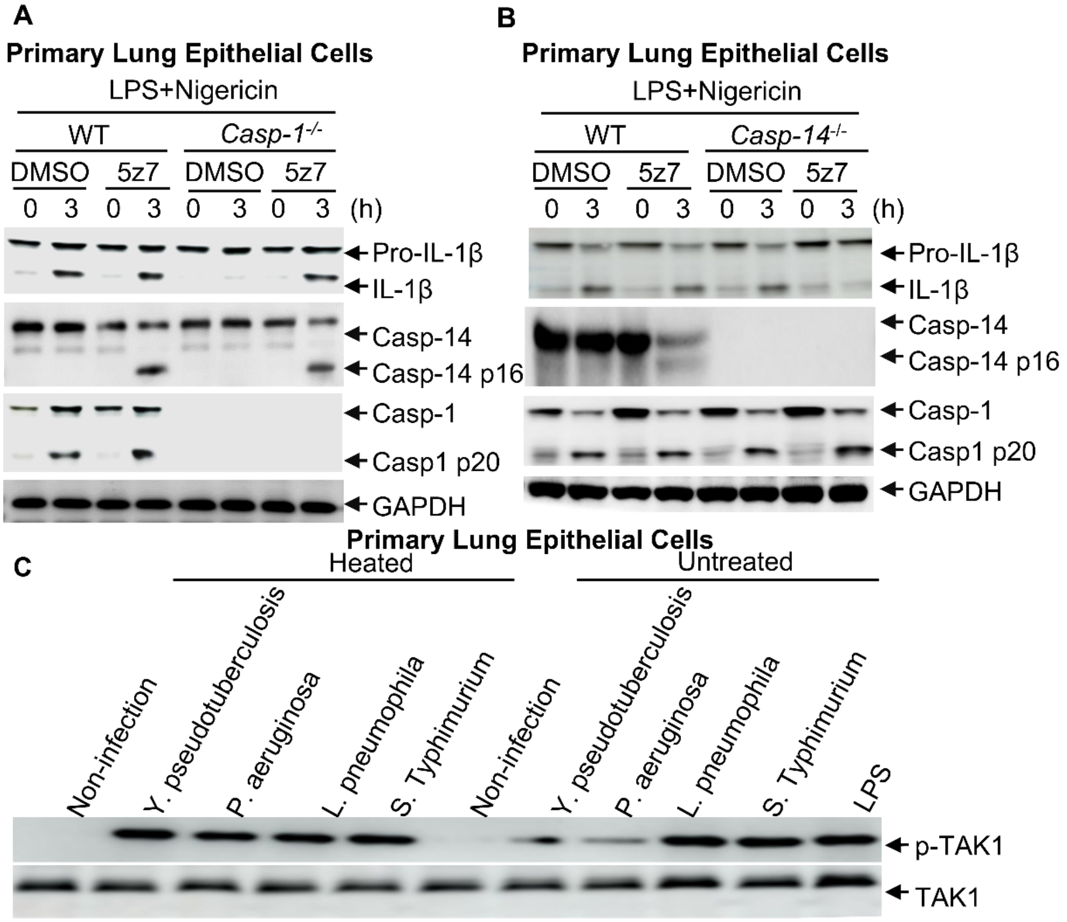


**Extended Data Fig. 4. TAK1 phosphorylates caspase-1 at S126**

**(A)**Immunoblotting of indicated proteins from epithelial cells of WT or *Caspase1*$^{-/-}$ *Caspase11*$^{-/-}$ stimulated with LPS/nigericin in the absence or presence of TAK1 inhibitor 5Z-7-Oxozeaenol for the indicated times. **(B)**Immunoblotting of indicated proteins from epithelial cells of *WT* or *Caspase14*$^{-/-}$ stimulated with LPS/nigericin in the absence or presence of TAK1 inhibitor 5Z-7-Oxozeaenol for the indicated times. **(C)** Immunoblotting of indicated proteins from epithelial cells with *Y. pseudotuberculosis, P.aeruginosa*, *L. pneumophila,* or *S. Typhimurium* infection MOI=5 for 12h. All of the immunoblot data are representative images from one of three independent experiments.

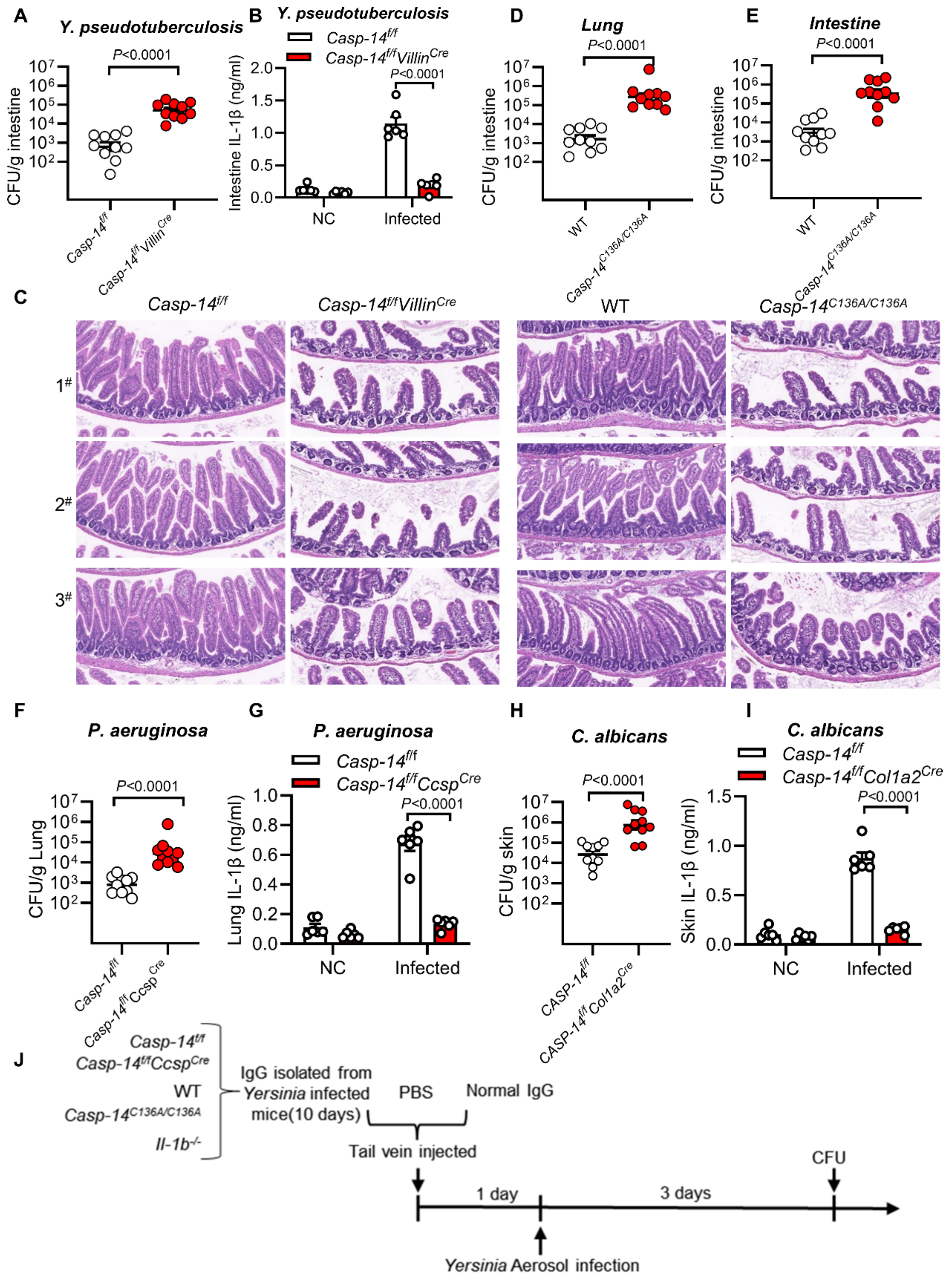


**Extended Data Fig. 5. Caspase-14-dependent maturation of IL-1β protects host against infection**

**(A-C)** 6-8 weeks female *Casp-14*$^{f/f}$ or *Casp-14*$^{f/f}$ *Villin*$^{cre}$ mice orally challenged with *Y. pseudotuberculosis* (~$10^7$ CFU/mice). CFU in intestine (a), IL-1β release serum (b) and H&E staining of lung tissue (c) (scale bar, 200 μm) of at 3 days post infection. **(D, E)** 6-8 weeks female *WT* or *Casp-14* $^{C132A}$ mice orally challenged with *Y. pseudotuberculosis* (~200 CFU/mice). CFU in lung/intesitine of at 3 days post infection.

**(F, G)** 6-8 weeks female *Casp-14*$^{f/f}$ or *Casp-14*$^{f/f}$ *Ccsp*$^{Cre}$ mice aerosol challenged with *P. aeruginosa* (~$10^6$ CFU/mice). CFU in lung (f) and IL-1β release serum (g) of at 3 days post infection. **(H, I)** 6-8 weeks female *Casp-14*$^{f/f}$ or *Casp-14*$^{f/f}$*KRT14*$^{Cre}$ mice skin challenged with *C. albicans* (~$10^6$ CFU/mice). CFU in lung (h) and IL-1β release serum (i) of at 3 days post infection. **(J)** Schematic diagram of the experimental process.

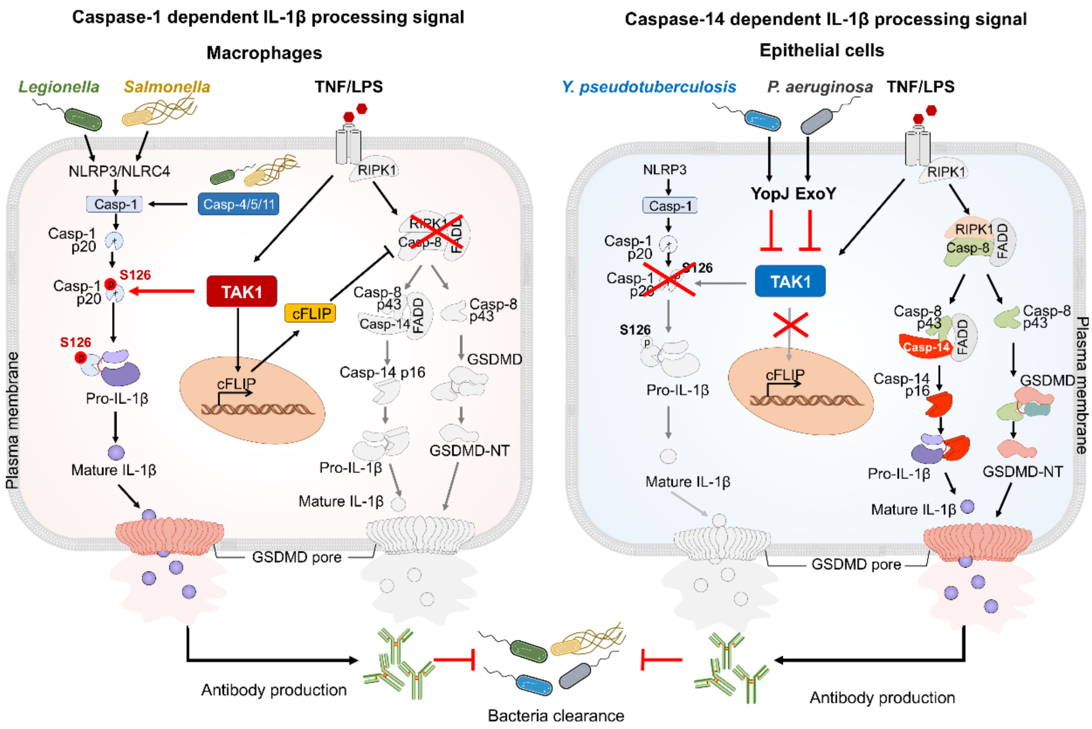


**Extended Data Fig. 6. Diagram**

In epithelial cells, pathogen blockade of TAK1 activation inhibits caspase-1-mediated cleavage of pro-IL-1β, thus leading to caspase-8–dependent cleavage of caspase-14, which directly interacts with and cleaves pro-IL-1β at Cys 132, thus leading to the inflammation and anti-*Y. pseudotuberculosis* humoral immunity.
